# Occupationally Relevant Wildfire Smoke Inhalation Impairs Nitric Oxide Signaling and Promotes Progressive Aortic Stiffening in Hypercholesterolemic Mice

**DOI:** 10.64898/2026.05.18.725908

**Authors:** Jacqueline Matz, Victoria A. Williams, Matthew J. Eden, Hannah I. B. Wilker, Simone Sabnis, Ye Chen, Paola Sebastiani, Michael J. Gollner, Jessica M. Oakes, Chiara Bellini

## Abstract

**Background:** Wildland firefighters experience repeated occupational exposure to wildfire smoke at high particulate matter (PM) concentrations, leading to elevated cardiovascular disease risk and hypertension prevalence. However, the pathophysiological processes linking cumulative smoke inhalation to vascular damage and blood pressure elevation remain poorly characterized. To evaluate these effects under controlled exposure conditions, we used a preclinical exposure model calibrated to match the cumulative PM burden deposited in wildland firefighter airways over 7-14 years of service. Male apolipoprotein E knockout (Apoe^−/−^) mice underwent whole-body inhalation of Douglas fir smoke or filtered air for 2 hours/day, 5 days/week, for 8 or 16 weeks at target PM concentrations of 40 mg/m^3^.

**Results:** Prolonged smoke exposure induced sustained elevation of circulating tumor necrosis factor-alpha (TNF-α), interleukin-1 beta (IL-1β), and interleukin-6 (IL-6), coupled with diffused nuclear factor kappa B (NF-κB) activation throughout the aortic wall. Smoke inhalation disrupted endothelial adherens junctions, upregulated intercellular adhesion molecule-1 (ICAM-1) and vascular cell adhesion molecule-1 (VCAM-1), and promoted monocyte recruitment to aortic tissues, concurrent with enhanced monocyte chemoattractant protein-1 (MCP-1) expression. Oxidative stress was evidenced by increased nicotinamide adenine dinucleotide phosphate (NADPH) oxidase subunit 2 (NOX2) expression, elevated superoxide levels, and endothelial nitric oxide synthase (eNOS) uncoupling in the aorta, leading to lipid peroxidation and accompanied by intimal apoptosis. These inflammatory and oxidative perturbations occurred alongside a pro-fibrotic phenotypic shift characterized by transforming growth factor beta 1 (TGF-β1) upregulation, myofibroblast differentiation, and progressive collagen accumulation in medial and adventitial compartments of the aortic wall. Functionally, smoke exposure progressively impaired aortic cyclic distensibility through combined wall thickening and circumferential tissue stiffening, while severely attenuating endothelium-dependent and nitric oxide (NO)-mediated vasodilation. These functional and structural shifts culminated in elevated systolic and diastolic blood pressures. While endothelial dysfunction reached maximal impairment by 8 weeks, aortic stiffening continued to worsen through 16 weeks of exposure, demonstrating differential temporal progression of vascular damage.

**Conclusions:** These findings demonstrate that occupationally relevant wildfire smoke exposure produces convergent inflammatory, oxidative, and profibrotic vascular remodeling with progressive loss of arterial compliance and impaired endothelium-dependent vasodilation, underscoring potential vascular targets for cardiovascular health surveillance and risk mitigation in wildland firefighters.

## INTRODUCTION

Wildfires are a leading source of episodic air pollution,^1^ generating emissions that can spread over large geographic regions and persist downwind of the fire perimeter.^2^ The projected increase in frequency and intensity of wildland fires over the next decade is expected to amplify the cumulative burden of wildfire smoke exposure across affected populations.^3^ Inhalation of wildfire smoke delivers a complex mixture of fine particulate matter (PM) with aerodynamic diameter ≤2.5µm (PM_2.5_), carbon monoxide (CO), polycyclic aromatic hydrocarbons, and other toxic constituents into the respiratory system, with downstream access to the systemic circulation. Exposure to pollutants in wildfire smoke has been associated with a range of adverse health outcomes,^4,5^ including cardiovascular events.^6,7^ Consistent with these observations, wildfires impose a substantial public health burden in the United States, with estimated health-related costs exceeding $130 billion annually.^8^

At the frontline of wildfire suppression, wildland firefighters (WLFF) experience repeated, high-intensity exposure due to close proximity to active fires and prolonged, physically demanding work with limited respiratory protection.^9^ These conditions place WLFF in a uniquely vulnerable occupational group in whom recurring smoke inhalation may confer elevated long-term cardiovascular risk. Repeated wildfire smoke exposure has been associated with increased incidence of cardiovascular morbidity and mortality in WLFF, including arrhythmias^10^ and acute coronary events.^11^ Positive associations between cumulative wildfire smoke inhalation and hypertension have been reported in both wildfire smoke-exposed communities and occupationally exposed populations.^10,12,13^ However, WLFF exhibit a higher prevalence of prehypertension and hypertension compared with the general population.^14^ Because hypertension is a well-established risk factor for cardiovascular disease, defining the mechanisms by which repeated wildfire smoke inhalation contributes to cardiovascular damage in this occupational group remains a priority.

While limited longitudinal data in WLFF preclude direct assessment of cumulative occupational risk, human studies have documented immediate cardiovascular and systemic responses elicited by wildfire smoke exposure during firefighting activities. Across a work shift, WLFF experience transient elevations in circulating pro-inflammatory mediators^15–19^ and markers of oxidative stress,^20–22^ along with short-term bursts in arterial stiffness.^22^ Although these responses may initially reflect adaptive host-defense mechanisms against airborne particulates or gaseous species in wildfire smoke, persistent or recurrent activation of inflammatory and oxidative pathways may become maladaptive, promoting endothelial injury and vascular remodeling that ultimately alter vascular tone and taper aortic compliance.^23–25^ Whether repeated wildfire smoke inhalation converts these acute, reversible responses^26^ into sustained pathological alterations of the vasculature that are capable of supporting chronically elevated blood pressure remains unresolved.

To address this question, we developed a preclinical exposure model designed to approximate the cumulative inhaled particulate burden experienced over a wildland firefighting tenure of tunable duration. Using dosimetry-based scaling, we established exposure conditions under which mice and WLFF accrue comparable cumulative deposited particulate mass in the lower respiratory tract, when normalized to lung surface area.^27,28^ To emulate wildfires in the Western United States, we sourced leaves and branches from Douglas fir trees as combustion fuel and subjected mice to whole-body smoke inhalation. Using this model, we showed that prolonged exposure to smoldering smoke induces early cardiopulmonary dysfunction, evidenced by increased airway resistance, diminished left ventricular ejection fraction, and a progressive, near-linear rise in blood pressure across the exposure period. These findings demonstrated that occupationally relevant wildfire smoke inhalation disrupts cardiovascular homeostasis in mice in a manner consistent with outcomes reported in WLFF. However, the molecular drivers and pathophysiological bases of these cardiovascular responses have yet to be defined.

Growing evidence indicates that inhalation of combustion-derived pollutants can initiate cardiovascular injury via circulating bioactive mediators. Because serum from mice exposed to wood smoke mixtures or individual smoke constituents impairs vasomotor function in the naïve mouse aorta *ex vivo*,^29,30^ we hypothesized that smoke inhalation would release systemic factors capable of perturbing vascular function and hemodynamic regulation.^7^ Positioned at the interface between circulating blood and the vessel wall, the vascular endothelium is an early target for these blood-borne mediators.^31,32^ Repeated endothelial exposure to inflammatory and oxidative signals during firefighting may therefore provide a mechanistic route through which acute responses compound over time, contributing to vascular dysfunction and hypertension risk in WLFF.

Building on this framework, in this study we examine whether repeated smoke exposure disrupts endothelial integrity and impairs nitric oxide (NO)-mediated vasodilatory signaling, and whether these luminal disturbances propagate into deeper layers of the arterial wall to induce maladaptive tissue remodeling. Using complementary molecular, histological, and biomechanical approaches, we assess the impact of cumulative fire smoke inhalation on aortic cyclic distensibility under physiological loading. Finally, by comparing multiple exposure durations, we evaluate whether vasomotor and arterial wall remodeling unfold along distinct temporal trajectories with continued exposure. Together, this work defines exposure-dependent vascular alterations that bridge acute physiological perturbations and longer-term dysfunction relevant to cardiovascular risk in WLFF.

## METHODS

A schematic of the study design, exposure paradigm, longitudinal assessments, and terminal vascular endpoints is shown in Figure 1.

**Figure 1.**
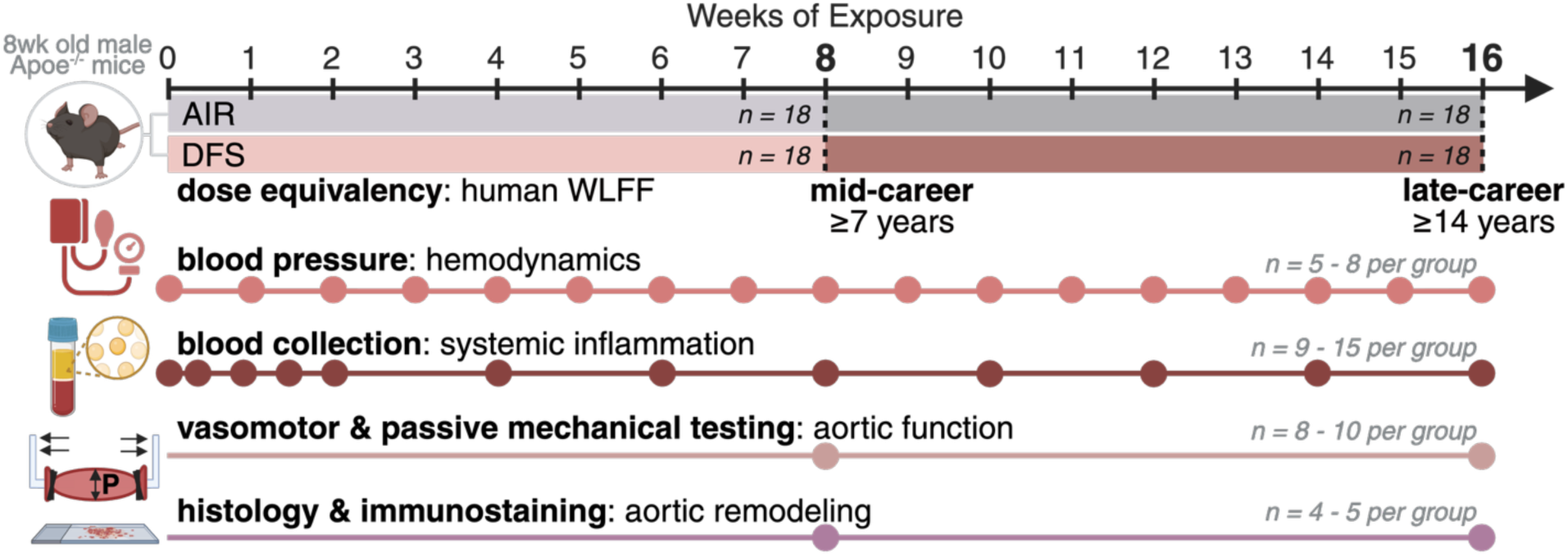
Overview of the experimental design used to investigate the vascular outcomes of occupationally relevant wildfire smoke exposure. Male Apoe^−/−^ mice were randomly assigned to whole-body inhalation of Douglas fir smoke (DFS) or filtered air (AIR) for 2 h/day, 5 days/week, for either 8 or 16 weeks, equivalent to cumulative exposures of 7+ and 14+ years of wildland firefighting service, respectively. Cardiovascular health and systemic inflammation were assessed longitudinally during the exposure period via noninvasive blood pressure measurements and serial blood collection for plasma cytokine analysis. At study endpoints, isolated suprarenal abdominal aortas were harvested for *ex vivo* assessment of vasomotor function and passive biaxial mechanical response. Parallel tissue cohorts were dedicated to histological, immunofluorescence, and *in situ* oxidative stress analyses to characterize aortic remodeling, inflammatory signaling, cell phenotype, and reactive oxygen species (ROS) generation.

### Mouse model of occupational wildfire smoke exposure

#### Animals

Male apolipoprotein E knockout (Apoe^−/−^) mice (N = 60) were obtained from The Jackson Laboratory (Bar Harbor, ME) at 8 ± 1 weeks of age. Male mice were used to reflect the predominantly male (>80%) WLFF workforce.^33^ Loss of apolipoprotein E impairs lipoprotein clearance, offering two key advantages for our application. First, Apoe^−/−^ mice develop chronic hypercholesterolemia, recapitulating elevated cholesterol levels reported in WLFF.^34^ Second, defective lipoprotein clearance creates a systemic pro-oxidative and pro-inflammatory environment. Because oxidative stress and inflammation were anticipated contributors to wildfire smoke-induced aortic stiffening and vasomotor dysfunction, the Apoe^−/−^ background provided enhanced sensitivity for detecting these vascular outcomes. Mice were housed under standard laboratory conditions with *ad libitum* access to standard chow and drinking water and maintained on a 12 h light-dark cycle. All animal procedures were approved by Northeastern University’s Institutional Animal Care and Use Committee (IACUC) and conducted in accordance with institutional guidelines.

#### Smoke exposure apparatus

Details of the custom smoke generation system and exposure setup have been reported elsewhere.^27^ Briefly, laboratory-scale wildfire smoke was produced from Douglas fir needles collected in Missoula, MT. Needles were separated from branches, oven-dried at 75°C for 72 h, and a dried fuel mass of 5.75 g was combusted under controlled smoldering conditions during two 1-h exposure sessions separated by a brief refueling interval. Smoke was generated using a ceramic heater operated at 450°C and mounted on a linear actuator that traversed a quartz tube packed with dried needles at 20 mm/min. Primary airflow was maintained at 3 L/min to sustain smoldering combustion, while secondary and tertiary airflow rates were set to 15 L/min and 4 L/min, respectively, to achieve target PM concentration of 40 mg/m^3^.

#### Exposure paradigm informed by dose equivalency

Using our cumulative exposure equivalency framework,^27^ 6,500 wildland firefighting hours (≥7 years of service depending on shift structure) mapped to 80 mouse exposure hours at the target PM concentration. Linear scaling of exposure equivalency with time yielded 160 mouse exposure hours for 13,000 hours (≥14 years) of service. To implement these exposures, an equal number of mice were randomly assigned to undergo whole-body inhalation of Douglas fir smoke (DFS) or filtered air (AIR, sham control) for 2 h/day, 5 days/week, for either 8 (8wk) or 16 (16wk) weeks. Smoke was delivered to mice in an exposure chamber following acclimation.^27^ Exposure chamber atmosphere was monitored daily by gravimetric measurement of PM using 47-mm glass fiber filter cassettes (Advantec, Dublin, CA) and continuous recording CO concentrations (Enerac 700; Enerac, Holbrook, NY). These data confirmed stable and reproducible exposure conditions, with PM concentrations (mean ± standard error of the mean) of 38.7 ± 0.8 mg/m^3^ and CO levels at 218.1 ± 6.5 ppm throughout the study.

#### Smoke exposure tolerability

To assess the potential for CO toxicity associated with DFS inhalation, blood carboxyhemoglobin concentrations were measured via submandibular venipuncture immediately following exposure at representative early, intermediate, and late time points (1 day, 8 weeks, and 16 weeks) using a blood gas CO-oximeter (ABL80 CO-OX OSM; Radiometer America, Brea, CA). Throughout the study, carboxyhemoglobin levels in DFS-exposed mice averaged 21 ± 4.7% and remained well below reported lethal thresholds for CO poisoning in mice (50-70%),^35^ while AIR-exposed controls consistently exhibited carboxyhemoglobin levels <3%. To evaluate overall systemic tolerability to DFS exposure, body mass was monitored weekly. Across the exposure period, body mass trajectories were comparable between DFS-and AIR-exposed mice, with similar weight gain over time.

### Longitudinal monitoring of cardiovascular health and systemic inflammation

#### Blood pressure measurement

Peripheral blood pressure was measured in awake, restrained mice (n = 8 per exposure group) using a noninvasive tail-cuff system (CODA; Kent Scientific, Torrington, CT). Measurements were obtained weekly and at least 18 h after the most recent exposure to minimize the influence of acute DFS inhalation on blood pressure.

#### Blood collection for detection of inflammatory markers

Whole blood samples (n = 15 per exposure group) were collected longitudinally via submandibular venipuncture into heparinized tubes at predefined time points: baseline; every other day between days 1 and 7; and every 2 weeks thereafter, corresponding to 0, 1, 3, 5, 7, 10, 20, 30, 40, 50, 60, 70, and 80 cumulative days of DFS or AIR exposure. Samples were centrifuged to isolate plasma, aliquoted, and stored at −80 °C until analysis. Systemic cytokine concentrations were quantified following a single freeze-thaw cycle using a commercial multiplex ELISA platform (Quansys Biosciences, Logan, UT) according to the manufacturer’s instructions.

### Assessment of isolated aortic function at study endpoints

#### Vasomotor function testing

Following either 8 or 16 weeks of exposure and 12-24 h after the final session, mice were deeply anesthetized with 2-3% isoflurane and euthanized by exsanguination. Whole aortas (n = 10 per exposure group and endpoint) were immediately excised and placed in cold (4 °C) Hank’s balanced salt solution (HBSS). Preparation of isolated aortic segments for vasoactive testing has been reported previously.^36^ Briefly, the suprarenal abdominal aorta was sectioned between the final pair of intercostal arteries and the renal arteries. Perivascular tissue was carefully removed, and lateral branches were ligated with single strands of braided 9-0 suture. Vessels were mounted on glass cannulae in a Krebs-Ringer solution bath and coupled to a custom computer-controlled biaxial testing system for simultaneous luminal pressurization and axial extension. The bath temperature was regulated at 37 °C, with continuous bubbling of 95% O_2_ and 5% CO_2_ to maintain physiological pH (7.4).

Following 10-min equilibration at preferred *in vivo* axial length and consistent luminal pressure of 80 mmHg, phenylephrine (2×10^-6^ M; L-Phenylephrine hydrochloride, 99%; Thermo Scientific, Waltham, MA) was added to the bath and aortic segments were allowed to reach stable vasoconstriction over 15 min. Endothelium-dependent vasodilation was evaluated by luminal administration of acetylcholine (acetylcholine chloride, 99%; Thermo Scientific) at increasing concentrations (10^-9^ to 10^-4^ M), separated by 5 min to achieve steady-state outer diameter. Upon completion of this challenge, vessels were returned to their unloaded configuration, and both the luminal HBSS and bath Krebs-Ringer solutions were replaced with fresh media. After 5-min equilibration, aortic segments were extended to their preferred *in vivo* configuration and incubated luminally with the nitric oxide synthase (NOS) inhibitor L-NAME (10^-4^ M; NG-Nitroarginine methyl ester hydrochloride; Tocris Bioscience, Bristol, UK) for 30 minutes. To probe the contribution of endothelial nitric oxide synthase (eNOS) to vasodilation, vessels were again pre-constricted with phenylephrine and subjected to a repeat acetylcholine dose-response protocol. Following a second washout and equilibration, endothelium-independent vasodilation was evaluated by luminal administration of the NO donor sodium nitroprusside (10^-9^ to 10^-4^; MP Biomedicals, Irvine, CA) after phenylephrine vasoconstriction. Throughout all protocols, outer diameter was continuously monitored using a camera positioned adjacent to the bath and interfaced with a custom LabVIEW-based acquisition system (National Instruments, Austin, TX). To account for variability in the degree of pre-constriction between aortic segments, vasodilation was expressed as percent reversal of phenylephrine-induced response and calculated as (*od*_dose_ − *od*_phen_)/(*od*_base_ − *od*_phen_) · 100, where *od*_dose_ is the steady-state outer diameter measured at each acetylcholine or sodium nitroprusside dose, *od*_phen_ is the outer diameter measured 15 min after phenylephrine administration and prior to vasodilator addition, and *od*_base_ is the baseline outer diameter measured before vasoconstriction.

#### Passive biaxial mechanical testing

At the conclusion of the vasomotor testing protocol, aortic segments (n = 10 per exposure group and endpoint) were returned to their unloaded configuration, and both the Krebs–Ringer bath and luminal HBSS were replaced with fresh HBSS. Passive biaxial testing protocols and data analysis have been described elsewhere.^36–38^ Briefly, vessels were equilibrated for 10 min at room temperature then acclimated and preconditioned under physiologically relevant biaxial loading. After estimation of unloaded geometry and determination of the preferred *in vivo* axial length, aortic segments were subjected to seven cyclic inflation-extension protocols: three pressure-diameter tests performed at, below, and above the preferred *in vivo* axial length, and four force-length tests conducted at luminal pressures of 10, 60, 100, and 140 mmHg. Following mechanical testing, cross-sectional rings were excised from the proximal end of each vessel and imaged under a dissection microscope to quantify unloaded wall thickness. Experimental data were fit by nonlinear regression to a microstructurally motivated four-fiber strain energy function of the form

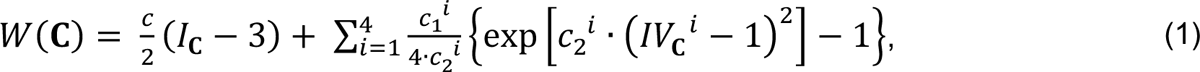

where *I*_*c*_ and *IV^i^_c_* are the first and fourth invariants of the right Cauchy-Green deformation tensor *c*. The first term represents the isotropic neo-Hookean contribution of elastic fibers and amorphous matrix, with parameter *c* having units of stress. The second term represents the anisotropic contributions of smooth muscle and collagen fiber families oriented along the axial (*i* = 1), circumferential (*i* = 2), and two symmetric diagonal (*i* = 3, 4) directions, with parameters *c^i^_1_* having units of stress and parameters *c^i^_2_* unitless. Best-fit material parameters were used to estimate structural (aortic geometry) and mechanical (aortic tissue-level) metrics at group-specific *in vivo* pressures. Linearized biaxial tissue stiffness about *in vivo* deformations was computed using small-on-large theory.^39^ Structural (geometry-dependent) stiffness of the aorta was quantified in terms of cyclic distensibility as (*id*_sys_ − *id*_dias_)/(*P*_sys_ − *P*_dias_), where *id* is the luminal diameter, *P* is the luminal pressure, and the subscripts *sys* and *dias* refer to systole and diastole, respectively.

### Tissue-level assessment of aortic remodeling and molecular signaling at study endpoints

#### Histological analysis of aortic wall composition

Following functional testing, a subset of aortic segments (n = 5 per exposure group and endpoint) was fixed in 10% neutral-buffered formalin for 18 h and transferred to 70% ethanol for storage. Fixed vessels were embedded in paraffin and serially sectioned at 5 μm thickness starting ∼500 μm from the proximal end, with three cross-sections mounted per slide. Aortic tissues were stained with either Movat’s pentachrome (MOVAT) or Picrosirius Red (PSR). MOVAT-stained cross-sections were imaged at 40x using a slide-scanning system (EasyScan Infinity; Motic, Kowloon Bay, Hong Kong). PSR-stained cross-sections were imaged at 40x under polarized light for visualization of birefringent collagen fibers using an upright microscope (DM4 B; Leica Biosystems, Wetzlar, Germany), with individual fields stitched to reconstruct complete cross-sections (Image Composite Editor; Microsoft, Redmond, WA). Area fractions of microstructural constituents were quantified from MOVAT-stained (elastin, vascular smooth muscle cell cytoplasm, and glycosaminoglycans) and PSR-stained (collagen) cross-sections using custom MATLAB routines (MathWorks, Natick, MA) based on hue-saturation-lightness thresholding, as previously reported.^36–38^

### Immunofluorescence-based analysis of vascular inflammation, oxidative stress, and cell phenotype

Suprarenal abdominal aorta samples from a separate cohort of mice (n = 4 per exposure group and endpoint) were embedded in optimal cutting temperature compound (OCT; 4585, Thermo Fisher Scientific, Waltham, MA), flash-frozen, and stored at −80 °C until processing. Aortic segments were cryosectioned beginning ∼200 μm from the proximal end into 6 μm-thick cross-sections. Tissues were fixed for 15 min in 4% paraformaldehyde, rinsed with 1x phosphate-buffered saline (PBS), then blocked and permeabilized for 15 min using 0.1% Tween-20 in PBS. Following permeabilization, cross-sections were blocked with 5% bovine serum albumin for 45 min, rinsed in PBS, and incubated for 1 h with primary antibodies against inflammatory mediators and associated signaling pathways, including nuclear factor kappa B (NF-κB) p65 (1:100; Abcam, ab16502), tumor necrosis factor-alpha (TNF-α; 1:300; Abcam, ab183218), interleukin-1 beta (IL-1β; 1:200; Proteintech, 16806-1-AP), and interleukin-6 (IL-6; 1:100; Abcam, ab179570); endothelial junctional and adhesion proteins, including vascular endothelial cadherin (VE-cadherin; 1:100; Abcam, ab33168), intercellular adhesion molecule-1 (ICAM-1; 1:75; Abcam, ab222736), vascular cell adhesion molecule-1 (VCAM-1; 1:100; Abcam, ab134047); macrophage and chemokine markers, including cluster of differentiation 68 (CD68; 1:50; Abcam, ab283654) and monocyte chemoattractant protein-1 (MCP-1; 1:60; Abcam, ab308522); oxidative and nitrosative stress markers, including nicotinamide adenine dinucleotide phosphate (NADPH) catalytic oxidase subunit 2 (NOX2; 1:100; Thermo Fisher Scientific, PA5-76034), 3-nitrotyrosine (3-NT; 1:100; Abcam, ab110282), 4-hydroxynonenal (4-HNE; 1:100; Invitrogen, MA5-27570), and superoxide dismutase 1 (SOD; 1:100, Abcam, ab13498); apoptotic signaling proteins, including cleaved caspase-3 (1:200; Cell Signaling Technology, 9664); profibrotic and vasoactive signaling proteins, including angiotensin II type 1 receptor (AT1R; 1:200; Proteintech, 25343-1-AP) and transforming growth factor beta 1 (TGF-β1; 1:75; Thermo Fisher Scientific, PA5-120329); and smooth muscle cell and extracellular matrix markers, including alpha-smooth muscle actin (ACTA2; 1:100; Proteintech, 14221-1-AP), myosin heavy chain 11 (MYH11; 1:100; Santa Cruz Biotechnology, sc-6956), and collagen I (1:200; Abcam, ab270993). Apoptotic nuclei were detected using a terminal deoxynucleotidyl transferase dUTP nick end labeling (TUNEL) assay (ApopTag Plus *In Situ* Apoptosis Fluorescein Detection Kit; Sigma-Aldrich, S7111). Following PBS rinses, cross-sections were incubated for 1 h with species-specific secondary antibodies, including donkey anti-rabbit Alexa Fluor 647 (1:500; Abcam, ab150075), goat anti-mouse Alexa Fluor 568 (1:500; Thermo Fisher Scientific, A-11004), and chicken anti-rat Alexa Fluor 647 (1:500; Thermo Fisher Scientific, A-21472). Nuclei were counterstained with 4′,6-diamidino-2-phenylindole (DAPI; 1:5000; Thermo Fisher Scientific, 62248) for 10 min. Three cross-sections per mouse were stained, and two regions per cross-section were imaged at 40x using a Zeiss LSM 800 confocal microscope with ZEN software (Carl Zeiss Microscopy, Jena, Germany). Acquisition parameters were identical across experimental groups for each stain. Images were processed in ImageJ (National Institutes of Health, Bethesda, MD). Where appropriate, the intimal, medial, and adventitial layers of the aortic wall were segmented as previously described.^36^ For all markers except NF-κB, results are reported as mean fluorescence intensity normalized to area for either the entire cross-section or individual aortic compartments. Unless otherwise stated, fluorescence intensity is quantified from 3 aortic cross-sections per mouse and 4 mice per exposure group at each endpoint. Individual datapoints show the mean of 2 square regions per cross-section.

To quantify NF-κB pathway activation within the aortic wall, we used immunostaining to quantify the nuclear translocation of the p65 subunit of NF-κB as previously described.^40^ Briefly, we created a nuclei mask by thresholding the DAPI stain and colocalized it onto a thresholded NF-κB p65 stain. We measured the average number of pixels in the thresholded NF-κB p65 stain inside the nuclei mask. Then, we created a new mask of each nucleus’s surrounding cytoplasm and measured the average number of pixels in the cytoplasm of the thresholded NF-κB p65 stain. In conclusion, we quantified the translocation of NF-κB p65 by normalizing the mean pixels inside the nucleus with the mean pixels in the surrounding cytoplasm.

#### In situ detection of vascular reactive oxygen species

Reactive oxygen species (ROS) were detected in situ by dihydroethidium (DHE) staining of mouse aortic tissues according to published protocols.^36,41^ DHE is a membrane-permeable fluorescent probe that undergoes intracellular oxidation through both nonspecific and superoxide-dependent pathways to generate nucleic acid-binding products emitting red fluorescence. Specimens for this assay were collected from the same aortic segments used for immunofluorescence and rinsed with ultrapure water to remove residual OCT ahead of staining. Tissue cross-sections were incubated with 10 μM DHE (D1168, Invitrogen) at 37 °C for 20 min. To assess the contribution of uncoupled eNOS to superoxide formation, matched aortic segments were pre-incubated with L-NAME (500 µM; Tocris Bioscience, Bristol, UK) at 37 °C for 30 min in preparation for DHE staining. We have previously established a positive control for DHE oxidation by inducing superoxide production in vascular tissues with menadione.^36^ Superoxide specificity was further evaluated by treating aortic tissue sections with the superoxide dismutase mimetic Mn(III) tetrakis(4-benzoic acid) porphyrin chloride (MnTBAP; 100 µM; Sigma-Aldrich, St. Louis, MO) prior to DHE staining as a negative control. Three cross-sections per mouse were stained, and two regions per cross-section were imaged at 40x using a Zeiss LSM 800 confocal microscope with ZEN software (Carl Zeiss Microscopy, Jena, Germany). Identical acquisition parameters were used across experimental groups for all DHE-stained cross-sections, and results are reported as mean fluorescence intensity normalized to area for individual aortic compartments.

### 2.9. Statistics

Sample size was estimated a priori using preliminary data and comparable studies to achieve 80% power for detecting exposure- and exposure duration-related differences with a minimum effect size of 0.1 (Cohen’s *d*) at a significance level of 0.05. The number of samples and mice included in each dataset is reported in the corresponding figure captions.

We present the data as mean ± standard error of the mean, unless we applied a data transformation to satisfy normality assumptions. We evaluate differences between groups (e.g., AIR8wk vs. DFS8wk) using linear regression model. We perform pairwise comparisons via linear contrasts in the model, with p-values calculated based on the associated t-tests for each contrast. We also adjusted p value using Benjamini–Hochberg false discovery rate (FDR) procedure where multiple comparisons are conducted. Differences were considered statistically significant if FDR < 0.10.

## RESULTS

### Systemic inflammatory signaling and diffuse NF-κB activation in the aorta

We evaluated circulating pro-inflammatory cytokines as potential systemic mediators of cardiovascular responses to wildfire smoke inhalation, given their documented involvement in vascular disease^32^ and prior associations with occupational exposure. Serum IL-6 concentrations in WLFF increase across a work shift;^15^ plasma levels of IL-6 and IL-1β in urban-area fire service instructors are positively related to the number of fire scenarios completed within a month;^42^ and circulating TNF-α in WLFF is elevated post-exposure compared to off-season, with strong loading in the late-season principal component analysis signature linked to cumulative PM_2.5_ exposure.^26,43^ We therefore measured concentrations of IL-6, IL-1β, and TNF-α in mouse plasma longitudinally throughout treatment and before daily exposure, with temporal resolution intended to capture potential lags between exposure and subsequent biological or functional responses. Plasma TNF-α levels transiently increased after one week of DFS exposure but returned to baseline by week two. At week twelve, plasma TNF-α and IL-1β levels were significantly higher in DFS-exposed mice than in filtered-air controls and remained elevated through study completion. Plasma IL-6 concentrations rose significantly by week sixteen in DFS-exposed mice relative to filtered-air controls, a delay consistent with IL-6 induction by TNF-α and IL-1β.^44^ These findings demonstrate an early, transient TNF-α response followed by sustained multi-cytokine elevation during prolonged exposure (Figure 2A). Vascular cells, including endothelial cells (EC), directly engage TNF-α and IL-1β through constitutively expressed membrane receptors^45,46^ but rely on IL-6 trans-signaling due to the absence of membrane-bound IL-6 receptor.^47^ Therefore, the induction off circulating TNF-α, IL-1β, and IL-6 in DFS-exposed mice is mechanistically poised to elicit endothelium-mediated vascular responses through their respective receptor pathways. Given the ability of TNF-α and IL-1β to activate NF-κB signaling cascades, we examined NF-κB as a central node for integrating cytokine responses.^48^ We performed immunostaining for the p65 subunit of NF-κB in mouse aortic tissue and quantified its nuclear translocation as a measure of pathway activation.^49^ We observed an increased p65 nuclear-to-cytoplasmic ratio in aortic EC after 8 weeks of DFS exposure compared with filtered-air controls (Figure 2B). At both the 8- and 16-week endpoints, this response extended to the medial and adventitial compartments, indicating propagation of inflammatory signaling throughout the aortic wall (Figure 2B).

**Figure 2.**
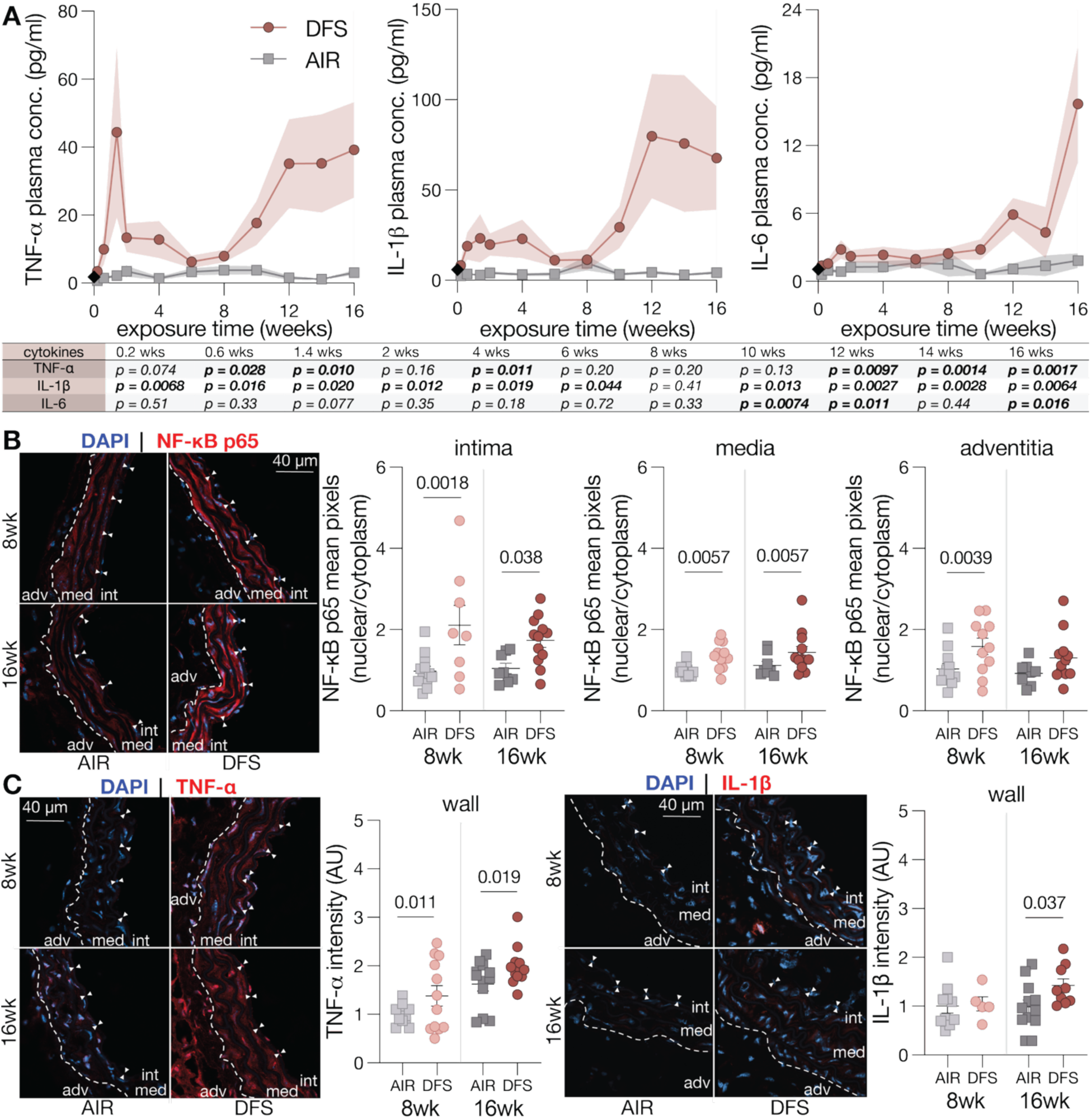
Nuclear factor kappa B (NF-κB) signaling in the mouse aorta associates with elevated circulating inflammatory mediators following wildfire smoke exposure. (**A**) Concentration of pro-inflammatory cytokines tumor necrosis factor-alpha (TNF-α), interleukin-1 beta (IL-1β), and interleukin-6 (IL-6) in mouse plasma over 16 weeks of AIR or DFS exposure. Datapoints show mean and SEM for up to 15 mice per exposure group at each endpoint. At day = 0, baseline from mice randomly chosen between groups is represented as black rhombus on graphs. (**B**) Representative immunofluorescence staining of NF-κB p65 (red) with 4′,6-diamidino-2-phenylindole (DAPI) nuclear counterstain (blue) in aortic tissues from AIR- and DFS-exposed mice. Quantification of nuclear p65 translocation based on p65/DAPI colocalization in the intima, media, and adventitia. (**C**) Representative immunofluorescence staining of TNF-α (red) with DAPI nuclear counterstain (blue) in aortic tissues from AIR- and DFS-exposed mice. (**D**) Representative immunofluorescence staining of IL-1β (red) with DAPI nuclear counterstain (blue) in aortic tissues from AIR-and DFS-exposed mice. Pairwise comparisons via linear contrasts in the linear regression model, with p-values calculated based on the associated t-tests for each contrast, with adjusted p-values to control for multiple comparisons in each panel. Log-transformed for panel (A). Statistical significance is indicated by p-values reported in the table or graphs for between-group differences.

NF-κB serves as a transcription factor regulating the expression of pro-inflammatory cytokines.^50^ TNF-α stimulation of murine vascular smooth muscle cells (VSMC) induces nuclear translocation of p65 and increases mRNA expression of inflammatory markers, including IL-1β and TNF-α;^51^ IL-1β positively autoregulates its own synthesis.^52^ We therefore quantified tissue accumulation of IL-1β and TNF-α by immunostaining. Consistent with the spatial distribution of NF-κB activation, expression of both cytokines extended across the aortic wall and was elevated in DFS-exposed mice compared with air-exposed controls (Figure 2C). Collectively, these findings demonstrate that DFS inhalation triggers systemic cytokine responses and drives diffuse aortic inflammation via NF-κB signaling.

### Disruption of endothelial adherens junctions, endothelial activation, and leukocyte recruitment to the aorta

Exposure to pro-inflammatory stimuli increases endothelial permeability to signaling molecules and induces the expression of adhesion receptors that facilitate immune cell recruitment to the vascular wall.^53^ TNF-α decreases membrane localization of VE-cadherin at adherens junctions in human umbilical vein EC and lung microvascular EC;^54,55^ IL-1β suppresses VE-cadherin transcription in lung microvascular EC;^56^ and IL-6 trans-signaling reduces VE-cadherin expression in human renal glomerular EC.^57^ Disruption of VE-cadherin mediated junctions is consistently associated with increased endothelial monolayer permeability.^58^ On this account, we evaluated VE-cadherin accumulation in the aortic endothelium and observed that DFS exposure diminished VE-cadherin fluorescence intensity relative to filtered-air controls at 16 weeks, consistent with compromised junctional integrity (Figure 3A). Because loss of barrier stability can facilitate inflammatory signaling, we next examined whether these structural changes were accompanied by upregulation of vascular adhesion molecules governed by NF-κB.^59^ Immunostaining showed increased ICAM-1 in the aortic endothelium of DFS-exposed mice compared with filtered-air controls at the 16-week endpoint, indicative of EC activation (Figure 3B). By contrast, VCAM-1 exhibited a transmural distribution across the aortic wall after 8 weeks of DFS exposure, reflecting a pro-inflammatory shift in resident vascular cell populations (Figure 3C).^60^

**Figure 3.**
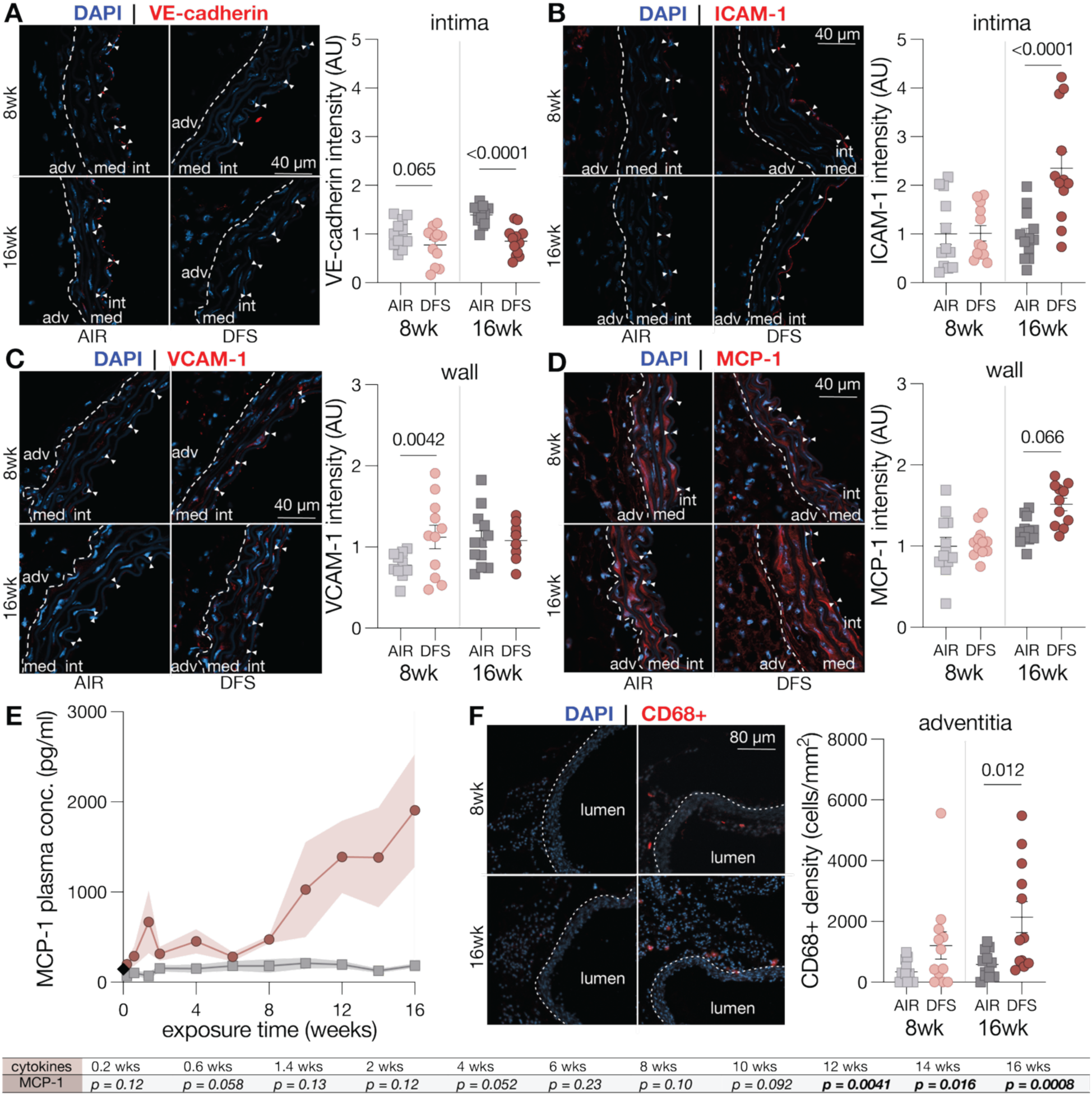
Leukocyte recruitment to the mouse aorta accompanies endothelial adherens junction disruption and endothelial activation following wildfire smoke exposure. (**A**) Representative immunofluorescent staining of vascular endothelial cadherin (VE-cadherin; red) with DAPI nuclear counterstain (blue) in aortic tissues from mice exposed to AIR or DFS. (**B**) Representative immunofluorescent staining of intercellular adhesion molecule-1 (ICAM-1; red) with DAPI nuclear counterstain (blue) in aortic tissues from AIR- and DFS-exposed mice. (**C**) Representative immunofluorescent staining of vascular cell adhesion molecule-1 (VCAM-1; red) with DAPI nuclear counterstain (blue) in aortic tissues from AIR- and DFS-exposed mice. (**D**) Representative immunofluorescent staining of monocyte chemoattractant protein-1 (MCP-1; red) with DAPI nuclear counterstain (blue) in aortic tissues from AIR- and DFS-exposed mice. (**E**) Concentration of pro-inflammatory chemokine MCP-1 in mouse plasma over 16 weeks of AIR or DFS exposure. Datapoints show mean and SEM for up to 11 mice per exposure group at each endpoint. At day = 0, baseline from mice randomly chosen between groups is represented as black rhombus on graph. (**F**) Representative immunofluorescent staining of cluster of differentiation 68-positive (CD68^+^) cells (red) with DAPI nuclear counterstain (blue) in aortic tissues from AIR- and DFS-exposed mice. Quantification of CD68^+^ cell density from 4 mice per exposure group at each endpoint and 3 aortic cross-sections per mouse. Plotted value from n=1 image of the entire aortic cross section of n=3 aortic rings for n=4 mice. Pairwise comparisons via linear contrasts in the linear regression model, with p-values calculated based on the associated t-tests for each contrast, with adjusted p-values to control for multiple comparisons in each panel. Log-transformed for panel (E). Statistical significance is indicated by p-values reported in the table or graphs for between-group differences.

Besides coordinating the expression of leukocyte adhesion proteins, NF-κB also induces the transcriptional upregulation of MCP-1, a key chemokine that directs leukocyte trafficking.^61^ Consistently, treatment with pro-inflammatory cytokines stimulates MCP-1 expression in vascular cells.^62^ We therefore evaluated MCP-1 expression within the aortic wall and observed greater fluorescence intensity in mice exposed to DFS for 16 weeks compared with air-exposed controls (Figure 3D). In addition, plasma MCP-1 levels were significantly higher in DFS-exposed mice than in controls from week 12 through the end of the study (Figure 3E). These results suggest that DFS inhalation generated chemokine gradients that could direct monocyte recruitment. To evaluate monocyte/macrophage infiltration in aortic tissues, we stained for the pan-macrophage antigen CD68 and observed that the density of CD68^+^ cells per aortic cross section was increased after 16 weeks of DFS exposure (Figure 3F), in line with recruitment mediated by MCP-1. Overall, these findings provide evidence that DFS inhalation disrupts endothelial barrier integrity, primes the endothelium for leukocyte adhesion, and establishes chemotactic signals that promote immune cell recruitment within the aortic wall.

### Oxidative and nitrosative stress, lipid peroxidation, and intimal apoptosis in the aorta

NF-κB activation drives transcription and expression of NOX2 in endothelial cells and fibroblasts of large arteries.^63,64^ Consistent with this mechanism, immunostaining revealed increased NOX2 signal in the aortic endothelium of mice exposed to DFS for 8 or 16 weeks, and in the aortic adventitia at the 16-week endpoint, compared to filtered-air controls (Figure 4A). NADPH oxidases serve as the principal source of superoxide in vascular cells,^65^ and endothelial overexpression of NOX2 increases vascular superoxide levels in Apoe^−/−^ mice.^66^ In addition to altered NOX expression, superoxide production is regulated by rapid post-translational activation of the enzyme complex.^65^ Pro-inflammatory cytokines act as potent stimuli that enhance NADPH oxidase activity and promote vascular reactive oxygen species (ROS) production.^65^ TNF-α induces oxidant release from human pulmonary artery and murine coronary microvascular EC through NADPH oxidase activation;^67,68^ NADPH oxidase mediates vesicular ROS generation in human aortic VSMC in response to TNF-α and IL-1β;^69^ IL-1β elicits superoxide formation in human umbilical vein EC and internal mammary artery VSMC;^70^ human dermal fibroblasts produce superoxide upon stimulation with IL-1β and TNF-α;^71^ Depending on the subcellular placement of NOX, superoxide and downstream ROS may localize within intracellular compartments or be released into the extracellular space.^72^ In vascular cells, NOX enzymes can generate superoxide that accumulates intracellularly and superoxide act as a trigger for cytosolic signaling pathways.^69,73,74^ On this basis, we assessed whether prolonged DFS inhalation altered the redox state of the mouse aorta. Aortic tissues from DFS-exposed mice exhibited significantly greater fluorescence of DHE oxidation products after 8 and 16 weeks compared with filtered-air controls (Figure 4B). Pre-incubation of DFS-exposed tissues with MnTBAP significantly reduced DHE fluorescence, confirming the selectivity of the signal for superoxide and serving as a negative control for the assay (Figure 4B). We have previously validated the DHE assay with additional controls, including naïve mouse aortic tissues treated with menadione as a positive control for superoxide generation, and pre-incubation with polyethylene glycol-conjugated superoxide dismutase (PEG-SOD) before menadione treatment or MnTBAP before DHE staining as additional negative controls.^36^

**Figure 4.**
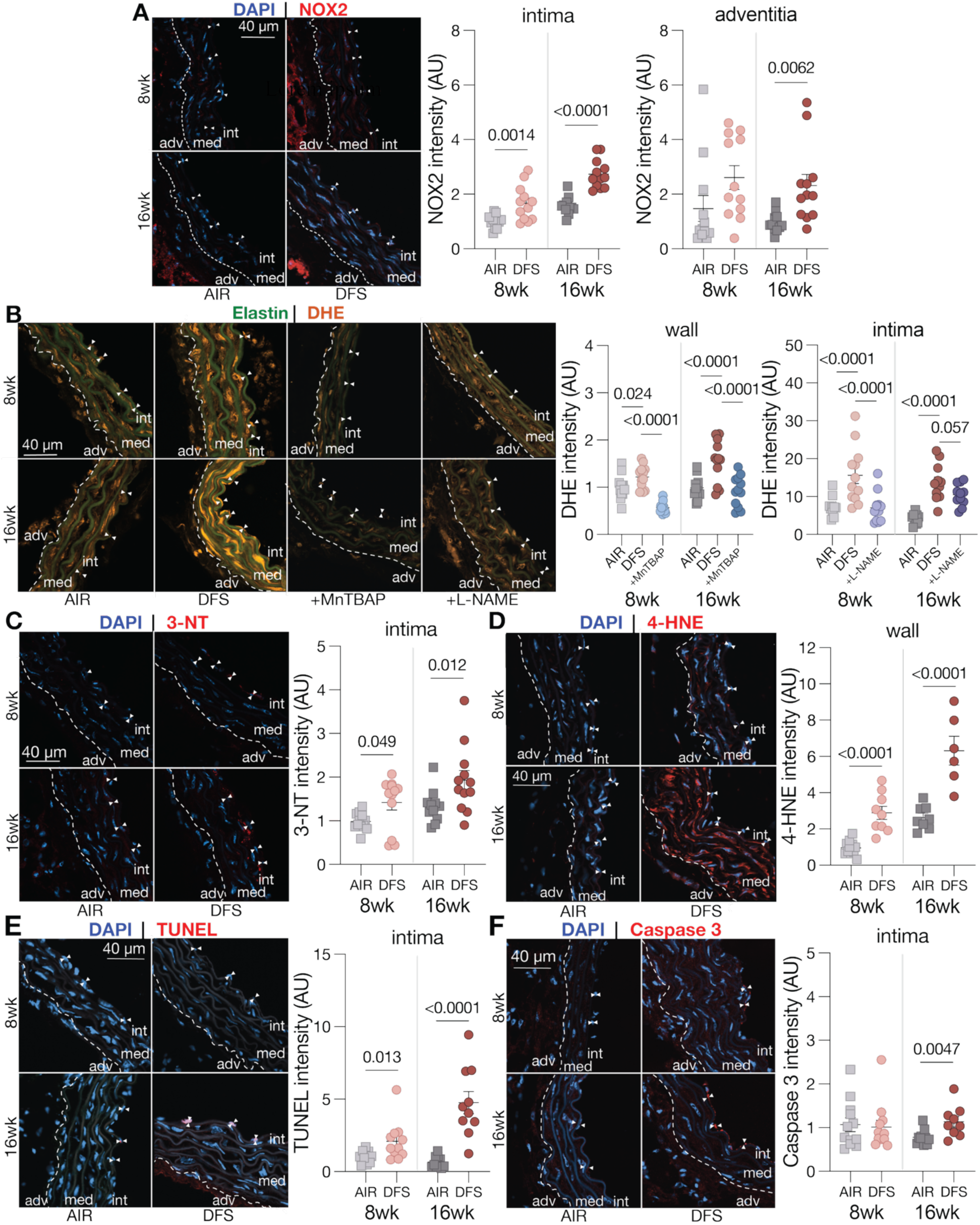
Endothelial cell apoptosis occurs alongside oxidative and nitrosative stress, lipid peroxidation, and endothelial nitric oxide synthase (eNOS) uncoupling in the mouse aorta following wildfire smoke exposure. (**A**) Representative immunofluorescent staining of nicotinamide adenine dinucleotide phosphate (NADPH) catalytic oxidase subunit 2 (NOX2; red) with DAPI nuclear counterstain (blue) in aortic tissues from mice exposed to AIR or DFS. (**B**) Representative fluorescence signal from dihydroethidium (DHE) oxidation products (orange) with elastin autofluorescence (green) in aortic tissues from AIR- and DFS-exposed mice. Tissue pre-incubation with the superoxide dismutase mimetic Mn(III) tetrakis(4-benzoic acid) porphyrin chloride (MnTBAP) and the nitric oxide synthase (NOS) inhibitor NG-nitroarginine methyl ester (L-NAME) was used to assess the specificity of the DHE signal for superoxide and the contribution of uncoupled eNOS to superoxide production, respectively. DHE oxidation products emit fluorescence in the red spectrum, but the signal is visualized in orange to enhance contrast. (**C**) Representative immunofluorescent staining of 3-nitrotyrosine (3-NT; red) with DAPI nuclear counterstain (blue) in aortic tissues from AIR- and DFS-exposed mice. (**D**) Representative immunofluorescent staining of 4-hydroxynonenal (4-HNE)-protein adducts (red) with DAPI nuclear counterstain (blue) in aortic tissues from AIR- and DFS-exposed mice. (**E**) Representative immunofluorescent staining of DNA fragmentation detected by terminal deoxynucleotidyl transferase dUTP nick end labeling (TUNEL) assay (red) with DAPI nuclear counterstain (blue) in aortic tissues from AIR- and DFS-exposed mice. (**F**) Representative immunofluorescent staining of cleaved (activated) caspase 3 (red) with DAPI nuclear counterstain (blue) in aortic tissues from AIR- and DFS-exposed mice. Pairwise comparisons via linear contrasts in the linear regression model, with p-values calculated based on the associated t-tests for each contrast, with adjusted p-values to control for multiple comparisons in each panel. Statistical significance is indicated by p-values reported in the table or graphs for between-group differences.

In addition to NOX, uncoupled eNOS constitutes another source of endothelial superoxide during vascular inflammation. Under physiological conditions, eNOS is constitutively expressed in EC to generate NO. Superoxide reacts with NO to form the reactive nitrogen species (RNS) peroxynitrite, which can promote eNOS uncoupling and divert the enzyme from producing NO to generating superoxide.^75^ To test whether uncoupled eNOS contributed to the oxidative burden in the endothelium of DFS-exposed mice, we incubated aortic tissues with the NOS inhibitor L-NAME prior to the DHE assay. This treatment significantly reduced intimal DHE fluorescence, implicating eNOS as a source of superoxide in this vascular compartment (Figure 4B). To gather additional evidence in support of eNOS uncoupling, we performed an indirect assessment of peroxynitrite activity by measuring 3-NT. Peroxynitrite nitrates protein tyrosine residues to form 3-NT, a stable end-product of nitrosative stress and an established marker of peroxynitrite-mediated protein modification.^76^ Immunostaining revealed increased accumulation of 3-NT-positive proteins along the luminal side of the aorta in DFS-exposed mice at the 8-week endpoint compared with filtered-air controls, with a similar trend observed at 16 weeks (Figure 4C).

ROS function as signaling mediators under physiological conditions but contribute to cellular injury and death when present at elevated levels due to either excessive production or impaired scavenging.^32,74^ To evaluate endogenous antioxidant capacity related to the latter, we quantified superoxide dismutase expression in aortic tissues and observed a significant decrease after 8 or 16 weeks of daily DFS inhalation compared with filtered-air controls (Figure S1). Given the oxidative and nitrosative stress detected in aortic tissues following DFS exposure, we next evaluated protein adducts formed through lipid peroxidation as adverse downstream molecular effects.^77^ Among the reactive aldehydes generated by this process, 4-HNE is a byproduct of polyunsaturated fatty acid peroxidation and an important mediator of oxidative protein modification.^78^ Immunostaining demonstrated greater levels of 4-HNE-protein adducts in aortic tissues of DFS-exposed mice at both 8 and 16 weeks, relative to filtered-air controls (Figure 4D). Similar to DHE oxidation products, 4-HNE was distributed throughout the aortic wall.

Excessive ROS and RNS can induce DNA damage that contributes to cell death. To determine whether these processes occurred in DFS-exposed mice, we first used the TUNEL assay to evaluate DNA fragmentation. DFS inhalation increased the number of TUNEL-positive nuclei in the aortic endothelium compared with filtered-air controls at both 8 and 16 weeks (Figure 4E). Because DNA fragmentation can also result from apoptosis in addition to direct oxidative injury, we next examined caspase-3 activation as a specific marker of apoptotic cell death. TNF-α initiates apoptosis in EC by recruiting caspase-8 to the receptor complex and triggering signaling cascades that culminate in the activation of effector caspase-3.^79,80^ Aligned with this mechanism, immunostaining revealed increased caspase-3 fluorescence intensity after 16 weeks of DFS exposure. Notably, both DNA damage and apoptosis markers were confined to the intima, despite oxidative stress and pro-inflammatory cytokine levels being elevated throughout the aortic wall (Figure 4F).

Collectively, these findings demonstrate that prolonged DFS inhalation induces persistent oxidative and nitrosative stress in the aorta, driven in part by uncoupled eNOS and enhanced NOX2 tissue expression, overwhelms antioxidant defenses, and promotes lipid peroxidation throughout the vessel wall, while DNA damage and apoptosis remain localized to the intima.

### IL-6 expression, pro-fibrotic phenotypic shift, and collagen-driven remodeling of the aorta

As second messenger species, intracellular ROS promote the release of additional pro-inflammatory mediators such as IL-6.^32,48^ ROS generated through NADPH oxidase activity in response to TNF-α or IL-1β mediate production of IL-6 and adhesion molecules via NF-κB signaling in human umbilical vein EC;^81–83^ activation of NF-κB by TNF-α and IL-1β requires ROS generated by NADPH oxidase in human aortic VSMC, and both cytokines stimulate IL-6 secretion in human VSMC.^69,84^ We therefore assessed IL-6 distribution in vascular tissues and observed increased expression throughout the aortic wall after 8 and 16 weeks of DFS exposure compared with filtered-air controls (Figure 5A). Because IL-6 upregulates AT1R transcription in rat aortic VSMC and murine aortic tissue,^85^ we examined whether DFS exposure promoted AT1R accumulation. The fluorescent signal for AT1R was broadly distributed across the vessel wall and was significantly elevated after 16 weeks of DFS exposure relative to filtered-air controls (Figure 5B). Since IL-6 treatment elicits diffuse superoxide generation in the mouse aorta independently of exogenous angiotensin II, and this effect is abolished in AT1R-deficient mice, our findings suggest that AT1R overexpression may further sustain vascular oxidative stress in response to DFS inhalation.^85^

**Figure 5.**
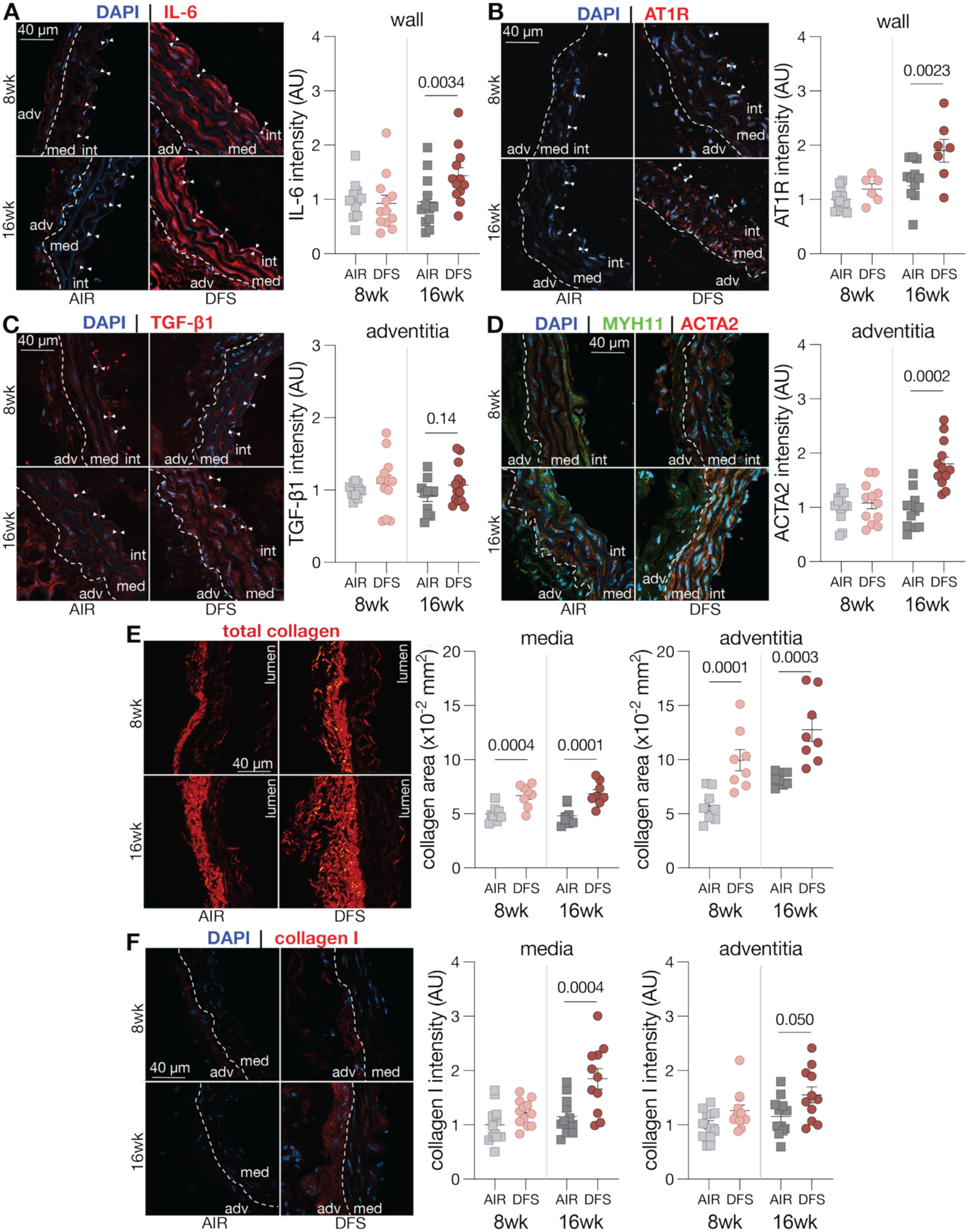
Pro-fibrotic phenotypic shift and collagen-driven remodeling in the mouse aorta align with inflammatory signaling enriched in IL-6 following wildfire smoke exposure. (**A**) Representative immunofluorescent staining of IL-6 (red) with DAPI nuclear counterstain (blue) in aortic tissues from mice exposed to AIR or DFS. (**B**) Representative immunofluorescent staining of angiotensin II type 1 receptor (AT1R; red) with DAPI nuclear counterstain (blue) in aortic tissues from AIR- and DFS-exposed mice. (**C**) Representative immunofluorescent staining of transforming growth factor beta 1 (TGF-β1; red) with DAPI nuclear counterstain (blue) in aortic tissues from AIR- and DFS-exposed mice. (**D**) Representative immunofluorescent staining of alpha-smooth muscle actin (ACTA2; red) and myosin heavy chain 11 (MYH11; green) with DAPI nuclear counterstain (blue) in aortic tissues from AIR- and DFS-exposed mice. (**E**) Representative picrosirius red staining of total collagen in aortic tissues from AIR- and DFS-exposed mice. (**F**) Representative immunofluorescent staining of collagen I (red) with DAPI nuclear counterstain (blue) in aortic tissues from AIR- and DFS-exposed mice. Pairwise comparisons via linear contrasts in the linear regression model, with p-values calculated based on the associated t-tests for each contrast, with adjusted p-values to control for multiple comparisons in each panel. Statistical significance is indicated by p-values reported in the table or graphs for between-group differences.

IL-6 signaling modulates the expression of TGF-β1 in fibroblasts and enhances TGF-β signaling in other cell types.^86,87^ Consistent with these reports, our analysis revealed a trend towards increased TGF-β1 accumulation in aortic tissues of DFS-exposed mice relative to filtered-air controls (Figure 5C). TGF-β1 is a pro-fibrotic cytokine that mediates collagen deposition and extracellular matrix remodeling through multiple mechanisms. Among these, IL-6-induced TGF-β1 production promotes the phenotypic transition of resident VSMC and fibroblasts into myofibroblasts with an enhanced capacity for collagen synthesis.^88,89^ Consistent with this process, immunostaining demonstrated elevated expression of ACTA2, a marker of fibroblast-to-myofibroblast differentiation, in the aortic adventitia following 16 weeks of DFS exposure compared with filtered-air controls (Figure 5D). These findings demonstrate that chronic DFS exposure promotes a pro-fibrotic shift within the vascular wall.

Given this evidence, we next examined whether molecular and cellular changes were accompanied by alterations in aortic geometry and microstructural organization. Morphometric analysis of MOVAT-stained cross-sections revealed increased total aortic cross-sectional area in DFS-exposed mice with respect to filtered-air controls, with significant enlargement of both medial and adventitial compartments (Table S1). This microstructural remodeling was not attributable to alterations in VSMC cytoplasm or elastin content, as both were preserved following DFS exposure, with elastin fibers maintaining their structural integrity (Figure S2). Instead, Picrosirius red staining revealed enhanced collagen accumulation in both medial and adventitial layers after 8 and 16 weeks of DFS exposure relative to filtered-air controls (Figure 5E). Immunostaining for collagen I, the predominant fibrillar collagen within the aortic wall, further corroborated such changes (Figure 5F). Collectively, these data establish that prolonged DFS inhalation promotes collagen-driven fibrotic remodeling of the aorta.

### Early circumferential tissue stiffening and progressive loss of cyclic distensibility in the aorta

Histological evidence of tissue fibrosis suggested that DFS inhalation may impair the ability of the aorta to function as a compliant pressure reservoir throughout the cardiac cycle. Given the prognostic significance of aortic stiffening in cardiovascular disease, we assessed the cyclic distensibility of the aorta in mice and quantified the contributing geometric and material determinants. Because hemodynamic loads directly influence vessel dimensions and tissue behavior, we first obtained blood pressure measurements to contextualize mechanical properties (Figure S3). Baseline data revealed no inter-group differences prior to exposure, with pooled values averaging 100 ± 1 mmHg systolic and 70 ± 1 mmHg diastolic across all animals. DFS inhalation increased blood pressure relative to controls, reaching on average 105 ± 1 mmHg at 8 weeks and 107 ± 1 mmHg at 16 weeks for systolic pressure, and 73 ± 1 mmHg at 8 weeks and 75 ± 2 mmHg at 16 weeks for diastolic pressure. Consistent with these shifts, pulse pressure was elevated in DFS-exposed mice after 16 weeks, with average values of 32 ± 1 mmHg compared with 26 ± 1 mmHg in controls. In contrast, mice exposed to filtered air maintained blood pressures near baseline throughout the study. To evaluate the effect of prolonged DFS exposure on geometry and tissue properties that govern the passive distension of the aorta under hemodynamic loads, excised vessels were subjected to controlled pressure cycles following restoration to their *in vivo* axial length. Consistent with histological findings, DFS inhalation significantly increased systolic wall thickness after both 8 and 16 weeks of exposure relative to filtered-air controls, with progressive thickening over time (Figure 6A). The structural remodeling of the aortic wall was accompanied by changes in circumferential tissue mechanics (Figure 6B). Mid-wall circumferential stretch at systolic pressure was elevated in the aorta of DFS-exposed mice at both endpoints with respect to filtered-air controls (Figure 6C), despite preserved luminal diameter. After 8 weeks of DFS inhalation, wall thickening offset the larger hemodynamic loads to maintain circumferential stress within the physiological range; after 16 weeks, however, continued remodeling overshot this effect, resulting in a trend towards attenuated wall stress compared with both age-matched, filtered-air controls and the 8-week DFS cohort (Figure 6C). Likewise, regulation of circumferential stress was contingent upon a concomitant increase in circumferential tissue stiffness, which was greater in DFS-exposed mice than in filtered-air controls at 8 weeks, while progressive wall thickening countered this effect at the 16-week endpoint (Figure 6C). The combination of elevated pulse pressure, wall thickening, and tissue stiffening led to a progressive loss of aortic cyclic distensibility, which worsened with longer DFS exposure and was more pronounced than in filtered-air controls (Figure 6D). DFS inhalation further impaired the axial recoil of the aorta, as evidenced by decreased axial stretch after 16 weeks of exposure compared with filtered-air controls (Figure 6E). Although this response contained axial wall stress and preserved axial stiffness (Figure 6E), it compromised aortic tissue capacity to store elastic energy in DFS-exposed mice at 16 weeks compared with age-matched filtered-air controls (Figure 6F). Collectively, these findings demonstrate that DFS inhalation augments both the intrinsic stiffness of aortic tissue and the structural stiffness of the aorta, thereby undermining its functional performance as a pressurized vessel designed to elastically distend and rebound under rapidly varying hemodynamic loads. Material parameters estimated from inflation-extension testing are reported in Table S2, with corresponding morphometric and mechanical metrics under simulated systolic conditions provided in Table S3.

**Figure 6.**
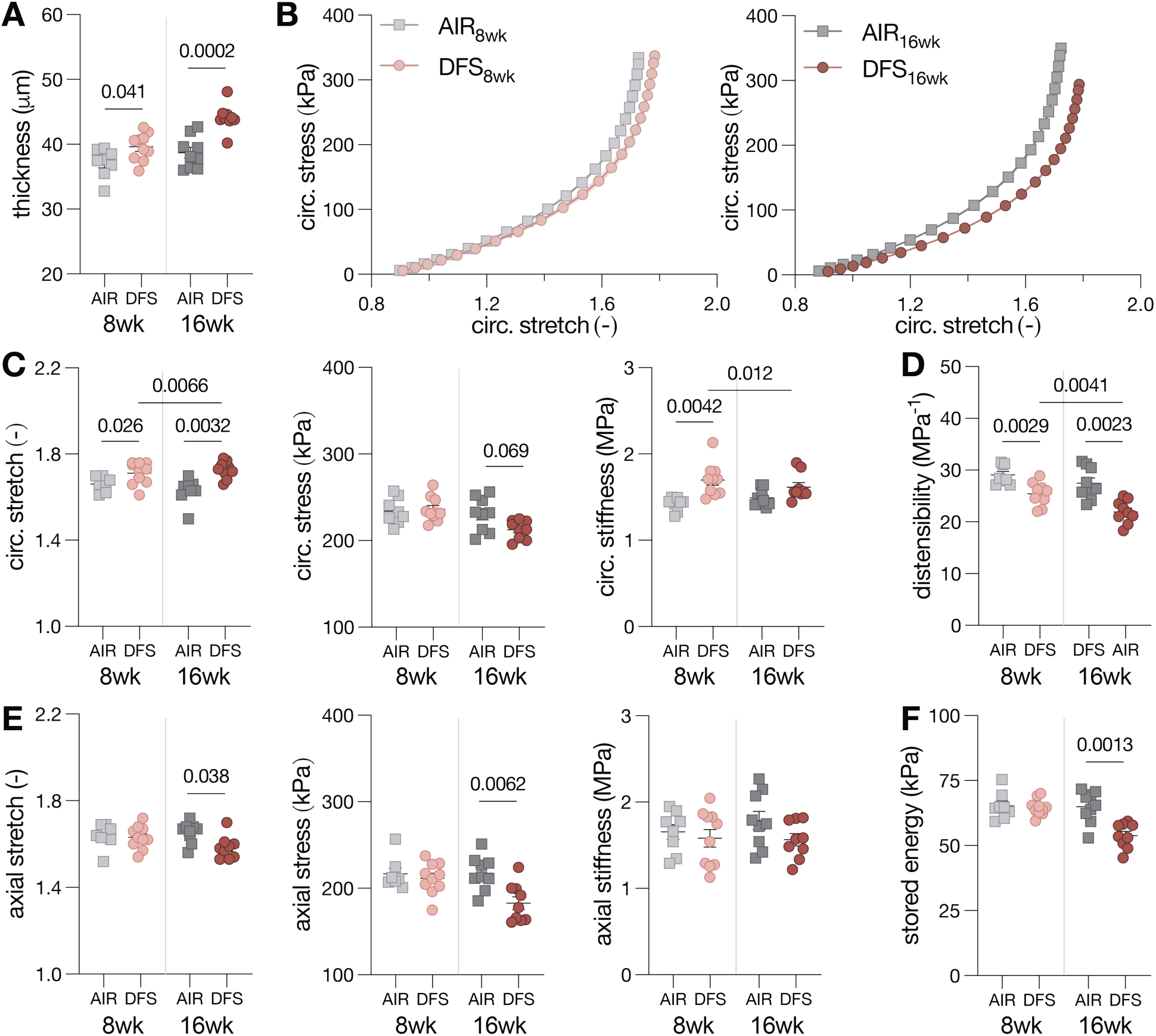
Structural stiffening of the mouse aorta following fire smoke inhalation. (**A**) Wall thickness of the aorta under *in vivo* biaxial loading in mice exposed to AIR or DFS. Datapoints represent measurements from individual mice with 10 mice per exposure group at each endpoint. (**B**) Circumferential mechanical behavior of aortic tissues from AIR- and DFS-exposed mice. Datapoints show mean with dispersion from 10 mice per exposure group at each endpoint. (**C**) Functional metrics of biaxial stretch, stress, and intrinsic tissue stiffness, together with structural cyclic distensibility and energy storage capacity of the aorta under *in vivo* loading. Datapoints represent measurements from individual mice with up to 10 mice per exposure group at each endpoint. Pairwise comparisons via linear contrasts in the linear regression model, with p-values calculated based on the associated t-tests for each contrast, with adjusted p-values to control for multiple comparisons in each panel. Statistical significance is indicated by p-values reported in the table or graphs for between-group differences.

### Disruption of NO-mediated vasodilation with preserved adrenergic vasoconstriction of the aorta

VSMC retain the capacity to respond to matrix stiffening by reorganizing their cytoskeleton to enhance actomyosin force production, a mechanism that could further limit aortic compliance *in vivo*.^90–92^ To probe this process in the context of DFS exposure, we examined contractile protein expression in aortic tissues and assessed vasomotor function in excised aortic samples. DFS inhalation increased ACTA2 accumulation in the aortic media while preserving MYH11 levels with respect to filtered-air controls at both endpoints (Figure 7A). However, vasoconstriction in response to consistent phenylephrine challenge was maintained in DFS-exposed mice relative to filtered-air controls at both 8 and 16 weeks (Figure 7B). As effectors of vascular tone, VSMC further mediate active vasodilation in part through the NO-soluble guanylyl cyclase-cyclic guanosine monophosphate signaling cascade that promotes smooth muscle relaxation. To assess whether this pathway remained functional following DFS exposure, we examined vasodilatory responses in excised aortic samples and found that the NO donor sodium nitroprusside elicited comparable dose-dependent relaxation across groups (Figure 7C). These data indicate that VSMC adrenergic contractility and relaxation mediated by exogenous NO were preserved following DFS inhalation. Nonetheless, the presence of nitrated tyrosine residues adjacent to the aortic intima and evidence of eNOS uncoupling indicate reduced bioavailability of endogenous, endothelium-derived NO in the aorta of DFS-exposed mice compared with filtered-air controls. Supporting this rationale, inflammatory mediators such as TNF-α suppress NO-mediated endothelium-dependent relaxation in mesenteric and coronary arteries.^93,94^ Guided by these observations, we investigated whether DFS exposure altered endothelium-dependent vasodilatory responses in aortic segments following cholinergic stimulation. DFS inhalation significantly attenuated acetylcholine-induced vasodilation in pre-constricted aortic segments relative to filtered-air controls following 8 and 16 weeks of exposure (Figure 7D). To assess whether impaired NO synthesis by eNOS contributed to this effect, aortic segments were pretreated with L-NAME prior to acetylcholine challenge. While eNOS inhibition further blunted vasodilation in control samples, no additional decline was observed in DFS-exposed mice at either endpoint (Figure 7D). Percent changes in diameter for these vasoactive challenges are summarized in Table S4. These data demonstrate that DFS inhalation disrupts endothelium-dependent dilation by impairing NO-mediated signaling.

**Figure 7.**
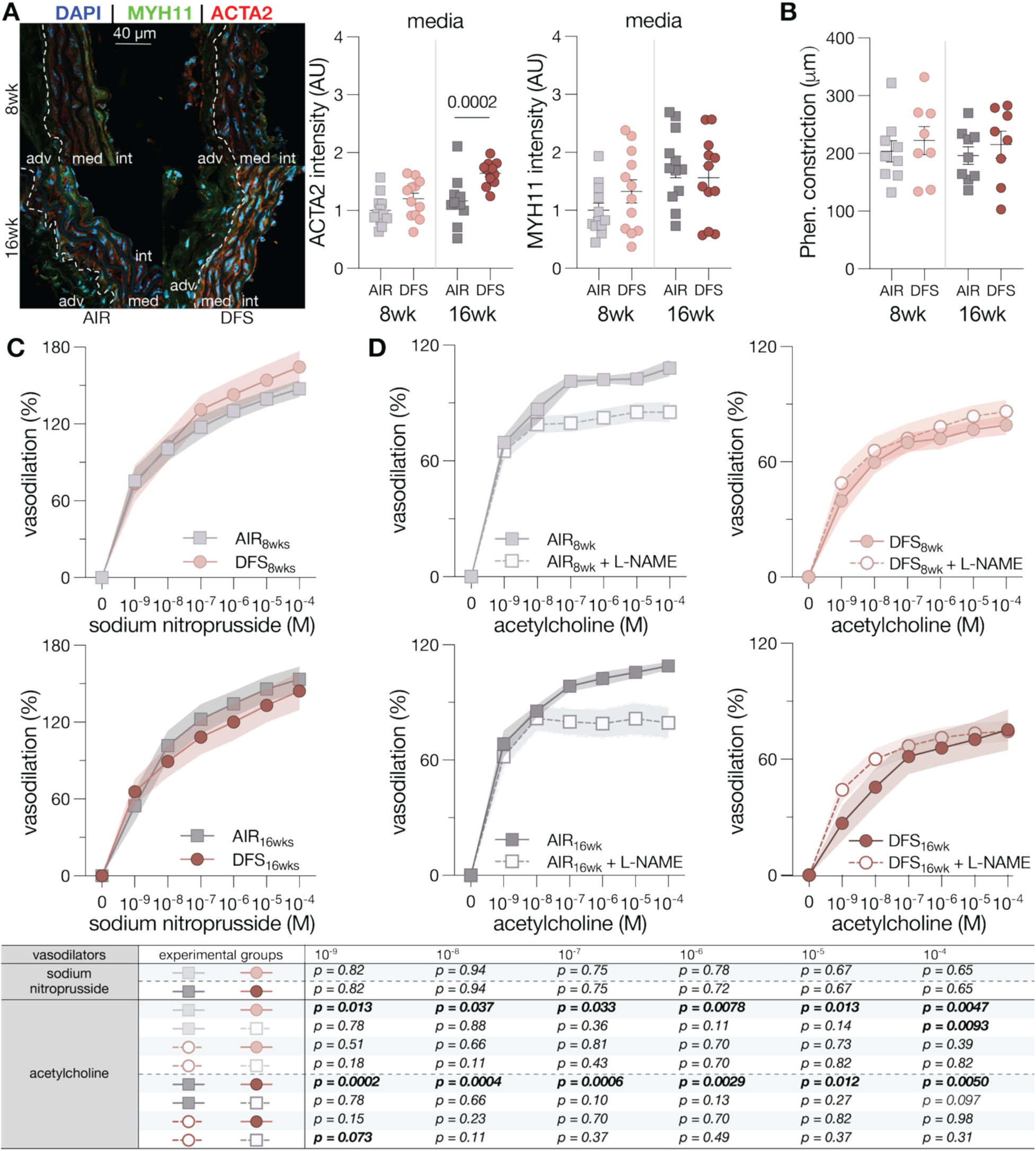
Impairment of endothelium-dependent and nitric oxide (NO)-mediated vasodilation in the mouse aorta following fire smoke inhalation. (**A**) Representative immunofluorescent staining of ACTA2 (red) and MYH11 (green) with DAPI nuclear counterstain (blue) in aortic tissues from mice exposed to AIR or DFS. **(B**) Maximum reduction in outer diameter of excised mouse aortic segments in response to phenylephrine (phen)-induced vasoconstriction. Datapoints show the mean of 3 phenylephrine challenges per mouse, with 8-10 mice per exposure group at each endpoint. (**C**) Endothelium-independent vasodilation as percent recovery of outer diameter from phenylephrine-induced vasoconstriction in response to increasing doses of the NO donor sodium nitroprusside. Datapoints show mean with dispersion for up to 10 mice per exposure group at each endpoint. (**D**) Endothelium-dependent vasodilation as percent recovery of outer diameter from phenylephrine-induced vasoconstriction in response to increasing acetylcholine doses. Pre-incubation of excised aortic segments with the NOS inhibitor L-NAME prior to phenylephrine challenge and acetylcholine stimulation were used to visualize the NO contribution to endothelium-dependent vasodilation. Datapoints show mean with dispersion from 8-10 mice per exposure group at each endpoint. Pairwise comparisons via linear contrasts in the linear regression model, with p-values calculated based on the associated t-tests for each contrast, with adjusted p-values to control for multiple comparisons in each panel. Statistical significance is indicated by p-values reported in the table or graphs for between-group differences.

Collectively, these findings suggest that DFS exposure may lead to an imbalance in vascular tone regulation.

## DISCUSSION

WLFF face recurrent physiological, environmental, and operational stressors that contribute to non-accidental occupational hazards.^95^ Comprehensive characterization of these risks is essential for informing effective mitigation strategies and supporting a sustainable, expanding workforce. Limited but growing evidence links wildland firefighting with a higher incidence of hypertension relative to the general population, with risk escalating as years of service accumulate.^10,96^ Our mouse model of wildfire smoke exposure, calibrated to match the cumulative particulate burden deposited in the lower respiratory tract of WLFF over the course of their career,^27^ recapitulates the rise in blood pressure and delineates two physiological mechanisms that mediate this response.

The first contributing process is the decline in aortic cyclic distensibility following repeated DFS inhalation. Such change reflects the structural stiffening of elastic arteries, which limits their ability to accommodate the stroke volume and results in larger systolic pressures consistent with those measured in DFS-exposed mice.^97^ Altered cyclic distensibility arises from both geometric remodeling and shifts in intrinsic tissue properties.^98^ The early increase in circumferential tissue stiffness is particularly noteworthy, as this parameter is tightly regulated through mechanosensing pathways responsive to extracellular matrix cues, thus even subtle elevations can meaningfully influence vascular mechanobiology.^99^ Additional mechanical impairments emerged over time, including reductions in axial stretch and stored energy, the latter of which is essential for normal aortic function because energy absorbed during systole facilitates elastic recoil and helps sustain diastolic perfusion.

The second precipitating event is the attenuation of endothelium-dependent vasodilation in response to acetylcholine, reflecting a disruption of NO-mediated signaling following long-term DFS inhalation. Although we did not directly assess smaller resistance vessels, the ubiquity of endothelial cells across the arterial system, together with evidence supporting a blood-borne source of injury following smoke inhalation, suggests that endothelial dysfunction may extend beyond the aorta to other vascular beds.^100^ Impaired endothelium-dependent vasodilation limits the ability of arteries to properly regulate vascular caliber in response to metabolic cues or shear stress.^101^ In contrast, DFS inhalation did not affect VSMC contractile function, as phenylephrine induced comparable vasoconstriction of the aorta across groups. Collectively, these findings support a shift in vasomotor balance toward a higher resting tone that elevates total peripheral resistance, consistent with the rise in diastolic pressure observed in DFS-exposed mice.^102^

By increasing the pressure against which the heart must eject blood (cardiac afterload), the combined effects of aortic structural stiffening and elevated peripheral resistance impose a greater mechanical burden on the left ventricle that may promote maladaptive cardiac remodeling over time.^24^ Notably, progressive loss of cyclic distensibility and endothelial dysfunction are hallmarks of aortic aging and serve as early determinants of future cardiovascular disease risk.^38^ In this context, our data suggest that prolonged inhalation of wildfire smoke at occupationally relevant concentrations encountered during wildland firefighting may accelerate cardiovascular aging, serving as a surrogate marker of biological aging.^98^

Consistent with features shared by many pathological remodeling processes, our findings implicate local oxidative and inflammatory signals as key mediators of DFS-induced aortic stiffening and altered vasomotor function. Plasma-derived factors appear to initiate these responses, which may then persist through feed-forward loops that translate systemic cues into sustained oxidative stress and inflammation across the aortic wall, establishing self-reinforcing processes that perpetuate vascular injury and adverse remodeling.

A first amplification loop involves direct NF-κB activation triggered by cytokine engagement of membrane receptors and unfolds through complementary mechanisms that reinforce systemic signals within aortic tissues. On the one hand, our data suggest that circulating TNF-α and IL-1β may stimulate endothelial cytokine production, renewing NF-κB-mediated inflammatory pathways over time and to neighboring cells, via autocrine and paracrine routes. On the other hand, our findings are consistent with increased endothelial permeability that may grant circulating inflammatory mediators access to deeper arterial layers, where they promote local cytokine production by intramural cells and extend the NF-κB response. Together, these processes establish a self-reinforcing inflammatory cycle that propagates throughout the aortic wall.

A second amplification loop is sustained by reciprocal interactions among cytokines, NF-κB signaling, and NADPH oxidase expression/activity across the aortic wall. Our data support coordinated regulation of the NOX pathway, whereby NF-κB-dependent transcription may enhance NOX expression, while TNF-α and IL-1β may directly promote NOX activation, collectively elevating vascular ROS levels. Notably, prior evidence demonstrates that ROS can then reinforce NF-κB activity: pharmacological inhibition of NADPH oxidase not only suppresses oxidant production, but also attenuates promoter transactivation downstream of NF-κB in EC stimulated with IL-1β, paralleling the effect of superoxide scavenging;^83^ NADPH oxidase subunit 1 (NOX1) and ROS modulate NF-κB engagement by IL-1β and TNF-α in VSMC;^69^ and oxidant signaling generated by NADPH oxidase contributes to NF-κB activation and the associated upregulation of adhesion molecules in EC stimulated by TNF-α.^103^ Our findings thus align with a feed-forward cycle in which cytokine-driven oxidative stress further amplifies NF-κB activity within aortic tissues.

A third amplification loop emerges through interactions among IL-6 signaling, angiotensin II pathways, and NADPH oxidase activity within the aortic wall. Our data suggest that IL-6-induced upregulation of AT1R may further potentiate NADPH oxidase-dependent ROS generation by enhancing the sensitivity of vascular cells to locally produced angiotensin II.^104^ Of note, deletion of the p65 subunit of NF-κB in mesenchymal cells protects against angiotensin II-induced expression of IL-6 and IL-1β.^105^ Therefore, our evidence implicates a self-sustaining cascade in which angiotensin II-mediated oxidative stress further reinforces NF-κB-driven inflammatory pathways throughout aortic tissues.

Lastly, the oxidative burden may have initially served as a trigger for, but was later reinforced by, eNOS uncoupling, contributing to sustained aortic inflammation in response to DFS exposure. Importantly, beyond these processes, the elevation in blood pressure driven by DFS-induced vascular stiffening and endothelial dysfunction can, in turn, aggravate oxidative stress, thereby compounding endothelial injury and fibrotic tissue remodeling.^106^

Our data further show that these feed-forward cycles unfolded over distinct timescales during repeated DFS inhalation. Reflecting this pattern, although oxidative stress and inflammation acted as a common initiating insult, downstream vascular outcomes evolved along different temporal trajectories. Endothelial dysfunction plateaued by 8 weeks of DFS inhalation, whereas aortic stiffening continued to worsen through the 16-week endpoint. Because the endothelium is directly exposed to circulating mediators, NO-dependent vasodilatory signaling was rapidly disrupted and stabilized at a relatively constant level of impairment. In contrast, progressive extracellular matrix remodeling within deeper wall layers supported the continued decline in cyclic distensibility with ongoing exposure. Collectively, these findings suggest that endothelial dysfunction emerges early due to direct luminal insult, while structural remodeling develops more gradually as damaging stimuli propagate into the arterial wall under the same upstream inflammatory-oxidative milieu.

Notwithstanding the many novel aspects of this work, some limitations warrant consideration. Although our data demonstrate clear increases in circulating pro-inflammatory mediators following DFS exposure, both the upstream stimuli that trigger their release and the cellular sources from which they originate remain uncertain. *In vitro* studies consistently demonstrate that respiratory cells are potent sources of cytokines and chemokines in response to wildfire smoke or constituents thereof. Wildfire PM upregulates transcription of genes encoding IL-1α, IL-1β, IL-8, and CCL20 in bronchial epithelial cells,^107^ and wildfire smoke extract elicits IL-6 release from differentiated primary bronchial epithelium.^108^ Biomass-smoke PM_2.5_ further activates both lung epithelial cells and alveolar macrophages, stimulating production of IL-17F along with downstream pro-inflammatory mediators such as IL-6, CCL2, and CCL4.^109^ Alveolar macrophages mount particularly robust responses: bushfire smoke extract increases intracellular active IL-1β and secreted IL-8,^110^ whereas PM_2.5_ from prescribed burns elevates TNF-α release,^111^ though responses depend on dosage.^112^ Cytokines generated in the lungs may enter the circulation and initiate broader systemic inflammatory responses. Additionally, peripheral blood mononuclear cells (PBMC) mobilized in response to pulmonary injury could further amplify cytokine production. This is supported by single-cell PBMC profiling of individuals exposed to wildfire smoke, including WLFF, which demonstrated that chronic exposure induces sustained epigenetic alterations related to cytokine signaling, thereby establishing a persistent pro-inflammatory environment capable of promoting vascular injury.^113^

The cytokine dynamics observed in our study are indicative of a biphasic inflammatory response. An initial, transient elevation in TNF-α is suggestive of an acute triggering event, whereas the prolonged increases in IL-6, IL-1β, and MCP-1 reflect sustained inflammatory signaling that develops with continued exposure. This pattern is consistent with prior evidence implicating early TNF-α activity in the initiation of endothelial activation and dysfunction.^114^ Defining how individual smoke constituents engage specific respiratory and immune cell populations to generate temporal cytokine patterns will be essential for elucidating how inhaled wildfire smoke initiates and sustains vascular inflammation. It also remains to be determined whether components of wildfire smoke that enter circulation further amplify these responses and contribute directly to cardiovascular injury. Furthermore, although our study used male mice to reflect the predominant composition of the firefighting workforce, it will be important to examine responses in females, who exhibit heightened sensitivity to specific smoke constituents.^115^ Finally, while our model isolates the effects of smoke exposure, concomitant environmental, physiological, and operational stressors in real-world firefighting scenarios may intensify adverse biological responses and should be considered when extrapolating laboratory findings to occupational settings.

## CONCLUSIONS

The prolonged and recurrent smoke exposure inherent to wildland firefighting underscores the need to understand the long-term cardiovascular consequences of this occupational hazard. In our preclinical cumulative exposure model, DFS inhalation increased circulating pro-inflammatory cytokines, activated the aortic endothelium, and produced diffuse inflammatory and oxidative perturbations throughout the vessel wall. Collagen-rich remodeling and increased aortic wall thickness accompanied these responses. This combined injury profile manifested functionally as progressive aortic stiffening and impaired endothelium-dependent vasodilation, leading to elevated systolic and diastolic pressures and reduced capacity to buffer pulsatile hemodynamic loads. These findings indicate that prolonged wildfire smoke inhalation can induce coordinated molecular, structural, and functional injury to the vasculature, raising important concerns regarding cardiovascular risk in WLFF.

## Abbreviations

PM: particulate matter
PM_2.5_: particulate matter with aerodynamic diameter ≤2.5µm
CO: carbon monoxide
WLFF: wildland firefighter
NO: nitric oxide
Apoe^−/−^: apolipoprotein E knockout
DFS: Douglas fir smoke
HBSS: Hank’s balanced salt solution
NOS: nitric oxide synthase
L-NAME: L-N-Nitroarginine methyl ester
eNOS: endothelial nitric oxide synthase
MOVAT: Movat’s pentachrome
PSR: Picrosirius Red
PBS: phosphate-buffered saline
NF-κB: nuclear factor kappa B
TNF-α: tumor necrosis factor-alpha
IL-1β: interleukin-1 beta
IL-6: interleukin-6
VE-cadherin: vascular endothelial cadherin
ICAM-1: intercellular adhesion molecule-1
VCAM-1: vascular cell adhesion molecule-1
CD68: cluster of differentiation 68
MCP-1: monocyte chemoattractant protein-1
NADPH: nicotinamide adenine dinucleotide phosphate
NOX2: NADPH catalytic oxidase subunit 2
NOX: NADPH catalytic oxidase subunit
3-NT: 3-nitrotyrosine
4-HNE: 4-hydroxynonenal
SOD: superoxide dismutase 1
AT1R: angiotensin II type 1 receptor
TGF-β1: transforming growth factor beta 1
ACTA2: alpha-smooth muscle actin
MYH11: myosin heavy chain 11
TUNEL: terminal deoxynucleotidyl transferase dUTP nick end labeling
DHE: dihydroethidium
MnTBAP: dismutase mimetic Mn(III) tetrakis(4-benzoic acid) porphyrin chloride
EC: endothelial cells
VSMC: vascular smooth muscle cells
ROS: reactive oxygen species
RNS: reactive nitrogen species
PBMC: peripheral blood mononuclear cells

## SUPPLEMENTARY INFORMATION

Supplementary material included as submission material.

## Acknowledgments

The authors acknowledge the Tufts Animal Histology Core (AHC) for histology services, the Institute for Chemical Imaging of Living Systems (RRID:SCR_022681) at Northeastern University for consultation and imaging support, and Ms. Hannah Kim for assistance with confocal image acquisition.

## Author contributions

JM: Methodology, Investigation, Formal Analysis, Data Curation, Visualization, Writing - Original Draft. VAW: Investigation, Formal Analysis, Visualization, Writing - Original Draft. MJE, HW, SS: Investigation, Writing - Review & Editing. YC, PS: Formal Analysis, Writing - Review & Editing. MJG, JMO: Conceptualization, Funding Acquisition, Writing - Review & Editing. CB: Conceptualization, Methodology, Supervision, Funding Acquisition, Writing - Original Draft.

## Data availability

The data that support the findings of this study are available from the corresponding author upon reasonable request.

## DECLARATIONS

### Ethics approval and consent to participate

All animal procedures were approved by the Institutional Animal Care and Use Committee (IACUC) at Northeastern University and were conducted in accordance with the Guide for the Care and Use of Laboratory Animals published by the U.S. National Institutes of Health.

### Consent for publication

Not applicable.

## SUPPLEMENTAL TABLES

**Table S1.** Microstructural composition of the aortic wall. Cross-sectional area of the aorta stratified by structural compartment and microstructural constituent in mice exposed to filtered air (AIR) or Douglas Fir smoke (DFS) for 8 or 16 weeks. Occasional small atherosclerotic lesions were excluded from analysis. Data are reported as mean ± SEM from 3 aortic cross-sections per mouse and 5 mice per exposure group at each endpoint. Pairwise comparisons were performed using t-tests with adjusted p-values to account for multiple comparisons for all panels. Statistical significance is indicated by p-values reported in the table or graphs for between-group differences. GAG: glycosaminoglycans.

| N = | AIR<sub>8wk</sub><br>5 | DFS<sub>8wk</sub><br>5 | AIR<sub>8wk</sub> VS.<br>DFS<sub>8wk</sub> | AIR<sub>16wk</sub><br>5 | DFS<sub>16wk</sub><br>5 | AIR<sub>16wk</sub> VS.<br>DFS<sub>16wk</sub> |
| --- | --- | --- | --- | --- | --- | --- |
| Total Area (x10<sup>-2</sup>mm<sup>2</sup>) |  |  |  |  |  |  |
| **Media** |  |  |  |  |  |  |
| Area | 12.53 ± 0.59 | 14.27 ± 0.36 | *p*=0.090 | 10.98 ± 0.30 | 13.98 ± 0.84 | ***p*=0.0069** |
| Elastin | 3.76 ± 0.19 | 3.33 ± 0.24 | *p*=0.29 | 2.63 ± 0.27 | 2.55 ± 0.22 | *p*=0.80 |
| Cytoplasm | 3.67 ± 0.28 | 4.02 ± 0.16 | ***p*=0.027** | 3.38 ± 0.57 | 4.41 ± 0.53 | ***p*=0.021** |
| Collagen | 4.95 ± 0.31 | 6.71 ± 0.52 | *p*=0.65 | 4.83 ± 0.40 | 6.88 ± 0.81 | *p*=0.42 |
| GAG | 0.14 ± 0.05 | 0.19 ± 0.05 | *p*=0.92 | 0.11 ± 0.04 | 0.12 ± 0.04 | *p*=0.93 |
| **Adventitia** |  |  |  |  |  |  |
| Area | 6.76 ± 0.79 | 11.32 ± 1.46 | ***p*=0.029** | 9.52 ± 0.36 | 14.09 ± 1.64 | ***p*=0.029** |
| Elastin | 0.22 ± 0.10 | 0.22 ± 0.07 | *p*=0.99 | 0.26 ± 0.06 | 0.33 ± 0.12 | *p*=0.99 |
| Cytoplasm | 0.67 ± 0.04 | 0.66 ± 0.07 | *p*=0.95 | 0.74 ± 0.07 | 0.70 ± 0.15 | *p*=0.95 |
| Collagen | 5.61 ± 0.68 | 10.16 ± 1.32 | ***p*=0.012** | 8.16 ± 0.28 | 12.78 ± 1.34 | ***p*=0.012** |
| GAG | 0.14 ± 0.01 | 0.23 ± 0.05 | *p*=0.29 | 0.25 ± 0.05 | 0.24 ± 0.05 | *p*=0.88 |
| **Wall** |  |  |  |  |  |  |
| Area | 19.29 ± 0.95 | 25.59 ± 1.33 | ***p*=0.001** | 20.50 ± 0.50 | 28.07 ± 1.17 | ***p*=0.0003** |
| Elastin | 3.98 ± 0.14 | 3.56 ± 0.28 | *p*=0.30 | 2.89 ± 0.31 | 2.87 ± 0.15 | *p*=0.97 |
| Cytoplasm | 3.67 ± 0.28 | 4.02 ± 0.16 | *p*=0.64 | 3.38 ± 0.57 | 4.41 ± 0.53 | *p*=0.42 |
| Collagen | 10.56 ± 0.91 | 16.87 ± 0.87 | ***p*=0.0001** | 13.01 ± 0.40 | 19.66 ± 0.99 | ***p*=0.0001** |
| GAG | 0.27 ± 0.05 | 0.41 ± 0.10 | *p*=0.77 | 0.36 ± 0.08 | 0.35 ± 0.06 | *p*=0.95 |

**Table S2.**
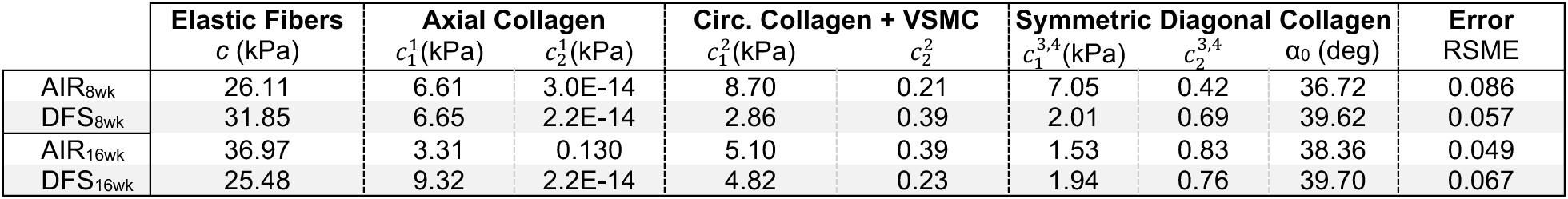
Constitutive descriptors of aortic tissue-level mechanical behavior. Best-fit material parameters estimated from averaged biaxial data using the four-fiber family constitutive model to describe the representative mechanical response of aortic tissues from mice exposed AIR or DFS for 8 or 16 weeks. VSMC: vascular smooth muscle cell.

**Table S3.**
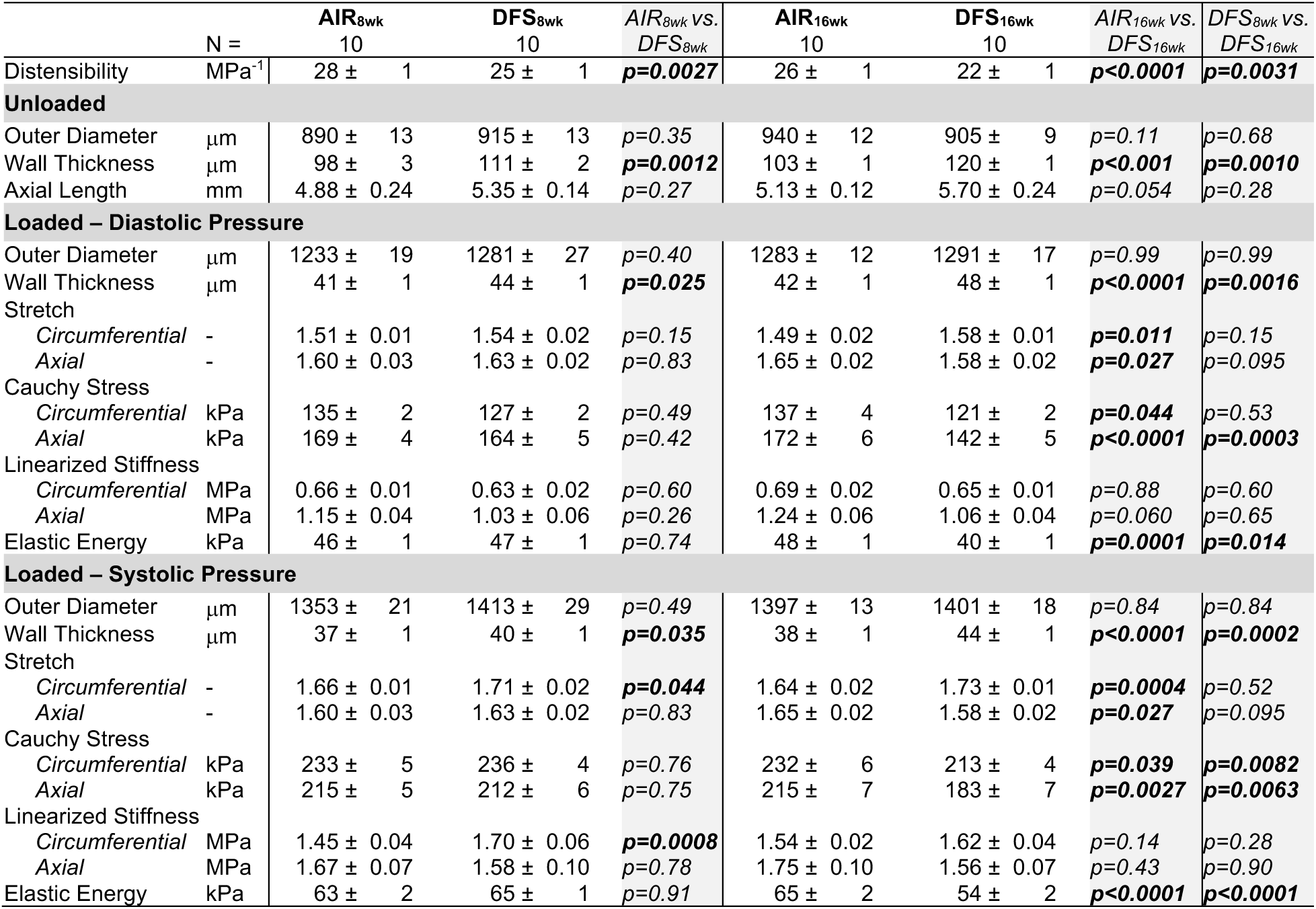
Structural and functional metrics of aortic mechanics under *in vivo* loading. Predicted structural and material properties of the aorta from mice exposed to AIR or DFS for 8 or 16 weeks. Data are reported as mean ± SEM from 10 mice per exposure group at each endpoint. Pairwise comparisons were performed using t-tests with adjusted p-values to account for multiple comparisons for all panels. Statistical significance is indicated by p-values reported in the table or graphs for between-group differences.

**Table S4.** Vasomotor function in excised aortic segments. Percent change in outer diameter in response to increasing concentrations of acetylcholine, acetylcholine following NG-Nitroarginine methyl ester hydrochloride (L-NAME) pre-treatment (10^-4^ M), and sodium nitroprusside. All vasoactive responses were assessed in aortic segments pre-constricted with phenylephrine (2·10^-6^ M). Ach: acetylcholine. SNP: sodium nitroprusside. Data are reported as mean ± SEM from 8-10 mice exposed to AIR or DFS for 8 or 16 weeks. Pairwise comparisons were performed using t-tests with adjusted p-values to account for multiple comparisons for all panels. Statistical significance is indicated by p-values reported in the table or graphs for between-group differences.

| 8 weeks | Naïve |  |  | L-NAME pretreatment |  |  |  |  |
| --- | --- | --- | --- | --- | --- | --- | --- | --- |
| Ach dose | AIR<sub>8wk</sub> | DFS<sub>8wk</sub> | AIR<sub>8wk</sub> vs. DFS<sub>8wk</sub> | AIR<sub>8wk</sub> | DFS<sub>8wk</sub> | AIR<sub>8wk</sub> vs. DFS<sub>8wk</sub> | AIR Ach vs. L-NAME + Ach | DFS Ach vs. L-NAME + Ach |
| $10^{-9}$ M | 69 ± 3 | 40 ± 3 | **$p = 0.013$** | 64 ± 2 | 49 ± 3 | $p = 0.18$ | $p = 0.78$ | $p = 0.51$ |
| $10^{-8}$ M | 86 ± 1 | 60 ± 2 | **$p = 0.037$** | 84 ± 2 | 66 ± 2 | $p = 0.11$ | $p = 0.88$ | $p = 0.66$ |
| $10^{-7}$ M | 96 ± 1 | 70 ± 2 | **$p = 0.033$** | 83 ± 2 | 72 ± 3 | $p = 0.43$ | $p = 0.36$ | $p = 0.81$ |
| $10^{-6}$ M | 102 ± 1 | 72 ± 2 | **$p = 0.0078$** | 83 ± 2 | 78 ± 2 | $p = 0.70$ | $p = 0.11$ | $p = 0.70$ |
| $10^{-5}$ M | 106 ± 1 | 77 ± 2 | **$p = 0.013$** | 87 ± 2 | 84 ± 2 | $p = 0.82$ | $p = 0.14$ | $p = 0.73$ |
| $10^{-4}$ M | 109 ± 1 | 76 ± 2 | **$p = 0.0047$** | 82 ± 3 | 86 ± 2 | $p = 0.82$ | **$p = 0.0093$** | $p = 0.39$ |
| 16 weeks | Naïve |  |  | L-NAME pretreatment |  |  |  |  |
| Ach dose | AIR<sub>16wk</sub> | DFS<sub>16wk</sub> | AIR<sub>16wk</sub> vs. DFS<sub>16wk</sub> | AIR<sub>16wk</sub> | DFS<sub>16wk</sub> | AIR<sub>16wk</sub> vs. DFS<sub>16wk</sub> | AIR Ach vs. L-NAME + Ach | DFS Ach vs. L-NAME + Ach |
| $10^{-9}$ M | 70 ± 2 | 65 ± 2 | **$p = 0.0002$** | 27 ± 2 | 44 ± 2 | **$p = 0.073$** | $p = 0.78$ | $p = 0.15$ |
| $10^{-8}$ M | 87 ± 3 | 79 ± 2 | **$p = 0.0004$** | 46 ± 3 | 60 ± 2 | $p = 0.11$ | $p = 0.66$ | $p = 0.23$ |
| $10^{-7}$ M | 101 ± 1 | 80 ± 2 | **$p = 0.0006$** | 61 ± 3 | 67 ± 2 | $p = 0.37$ | $p = 0.10$ | $p = 0.70$ |
| $10^{-6}$ M | 102 ± 1 | 83 ± 2 | **$p = 0.0029$** | 66 ± 3 | 71 ± 2 | $p = 0.49$ | $p = 0.13$ | $p = 0.70$ |
| $10^{-5}$ M | 103 ± 1 | 86 ± 2 | **$p = 0.012$** | 70 ± 3 | 73 ± 2 | $p = 0.37$ | $p = 0.27$ | $p = 0.82$ |
| $10^{-4}$ M | 108 ± 2 | 87 ± 2 | **$p = 0.0050$** | 75 ± 4 | 74 ± 2 | $p = 0.31$ | **$p = 0.097$** | $p = 0.98$ |

| 8 weeks | Naïve |  |  | 16 weeks | Naïve |  |  |
| --- | --- | --- | --- | --- | --- | --- | --- |
| SNP dose | AIR<sub>8wk</sub> | DFS<sub>8wk</sub> | AIR<sub>8wk</sub> vs. DFS<sub>8wk</sub> | SNP dose | AIR<sub>16wk</sub> | DFS<sub>16wk</sub> | AIR<sub>16wk</sub> vs. DFS<sub>16wk</sub> |
| $10^{-9}$ M | 72 ± 3 | 75 ± 3 | $p=0.82$ | $10^{-9}$ M | 66 ± 4 | 55 ± 5 | $p=0.82$ |
| $10^{-8}$ M | 95 ± 3 | 101 ± 3 | $p=0.94$ | $10^{-8}$ M | 89 ± 4 | 102 ± 4 | $p=0.94$ |
| $10^{-7}$ M | 124 ± 3 | 117 ± 3 | $p=0.75$ | $10^{-7}$ M | 108 ± 5 | 122 ± 5 | $p=0.75$ |
| $10^{-6}$ M | 135 ± 3 | 130 ± 2 | $p=0.78$ | $10^{-6}$ M | 120 ± 5 | 134 ± 4 | $p=0.72$ |
| $10^{-5}$ M | 147 ± 4 | 140 ± 2 | $p=0.67$ | $10^{-5}$ M | 133 ± 5 | 146 ± 4 | $p=0.67$ |
| $10^{-4}$ M | 157 ± 4 | 148 ± 2 | $p=0.65$ | $10^{-4}$ M | 144 ± 5 | 154 ± 3 | $p=0.65$ |

## SUPPLEMENTAL FIGURES

**Figure S1.**
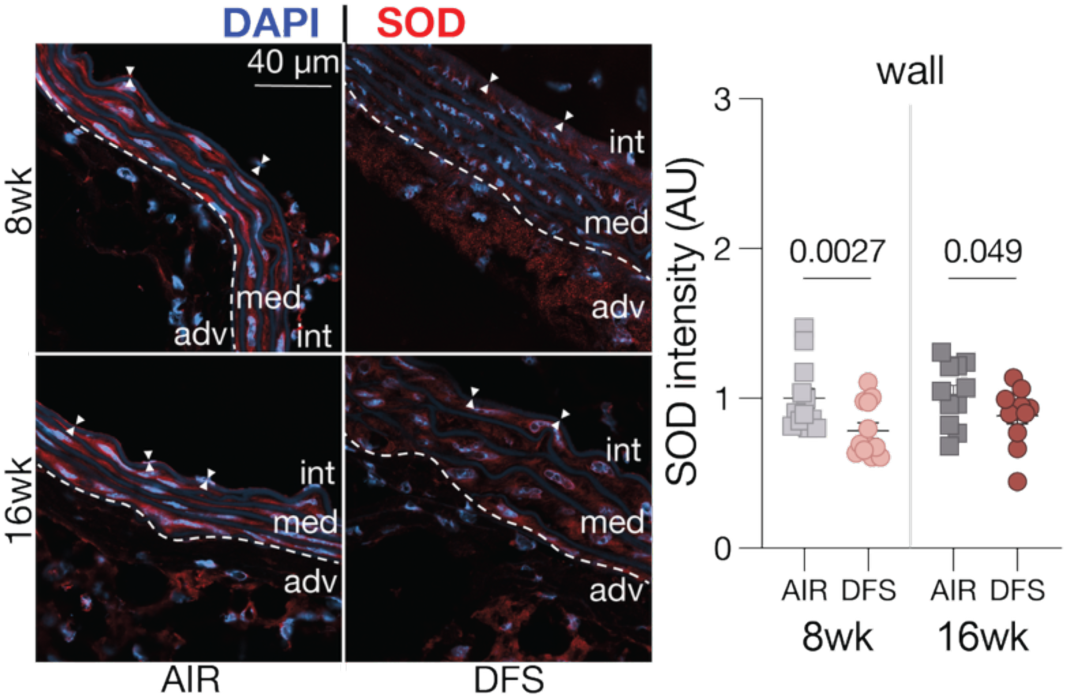
Attenuated endogenous antioxidant defenses in the mouse aorta following wildfire smoke inhalation. Representative immunofluorescence staining of endogenous superoxide dismutase (SOD; red) with 4′,6-diamidino-2-phenylindole (DAPI) nuclear counterstain (blue) in aortic tissues from mice exposed to filtered air (AIR) or Douglas fir smoke (DFS) for 8 or 16 weeks. Quantification of SOD fluorescence intensity through the wall from 3 aortic cross-sections per mouse and 4 mice per exposure group at each endpoint. Individual datapoints show the mean of 2 square regions per cross-section. Pairwise comparisons were performed using t-tests with adjusted p-values to account for multiple comparisons for all panels. Statistical significance is indicated by p-values reported in the graphs for between-group differences.

**Figure S2.**
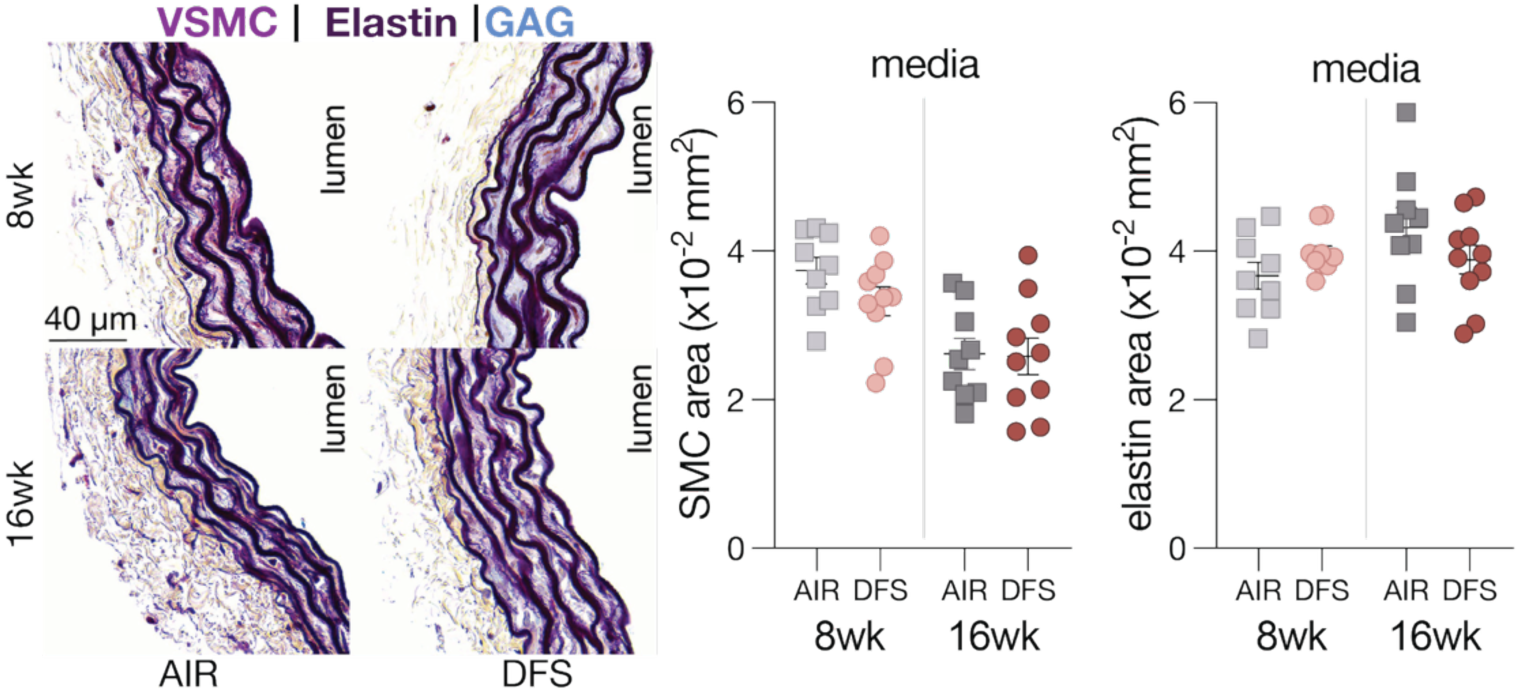
Preserved medial elastin and vascular smooth muscle cell (VSMC) content in the mouse aorta following wildfire smoke inhalation. (**A**) Representative Movat’s pentachrome (MOVAT) staining of elastin (dark purple and black) and VSMC cytoplasm (light purple) in aortic tissues from AIR- and DFS-exposed mice. Quantification of total elastin and VSMC cytoplasm content by area within the media averaged from 3 aortic cross-sections per mouse and 5 mice per exposure group at each endpoint. Pairwise comparisons were performed using t-tests with adjusted p-values to account for multiple comparisons for all panels. Statistical significance is indicated by p-values reported in the graphs for between-group differences. GAG: glycosaminoglycans.

**Figure S3.**
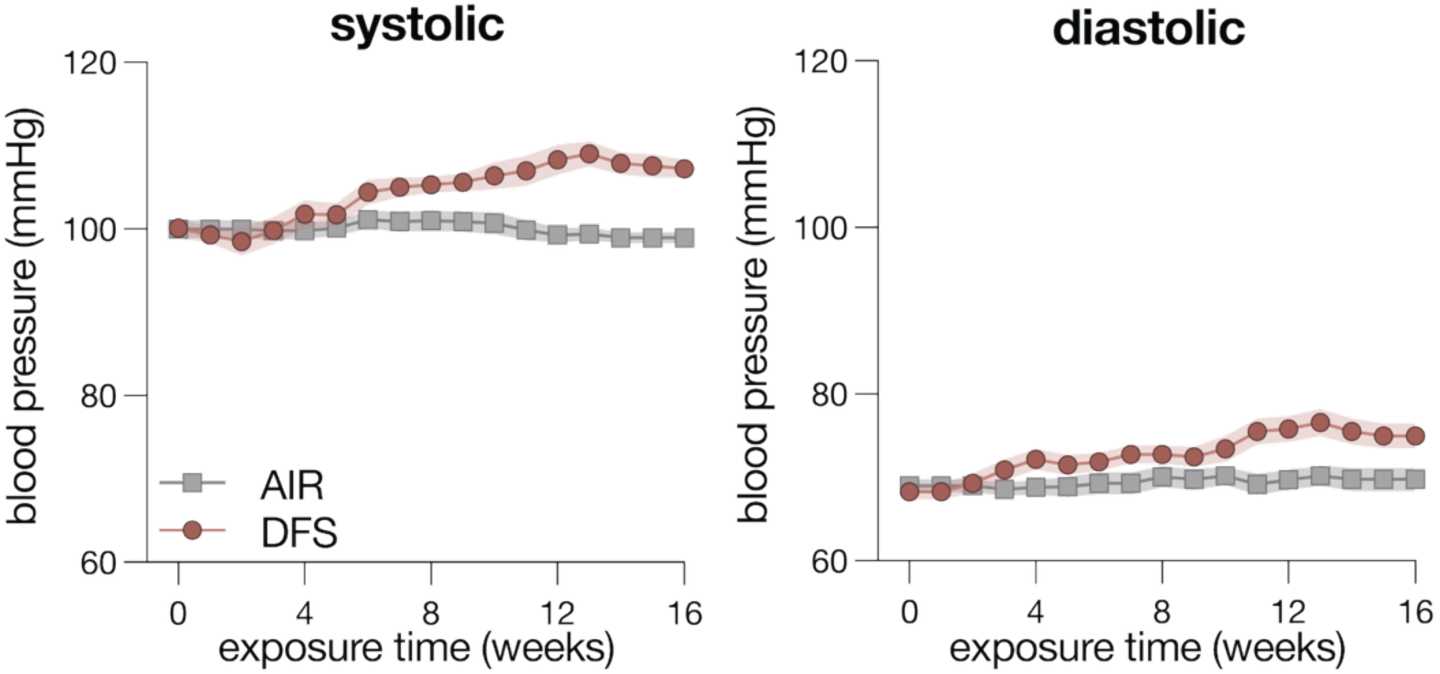
Elevated blood pressure in mice following wildfire smoke inhalation. Average systolic and diastolic blood pressure measured in mice exposed to AIR or DFS over the course of the study. Symbols represent weekly mean values from n = 8 mice per exposure group.

